# Live single-molecule imaging reveals global shifts in mRNA mobility during human stem cell differentiation

**DOI:** 10.64898/2026.07.27.740939

**Authors:** Aileen-Diane Bamford, Gilles Gut, Tim-Oliver Buchholz, Ryoko Okamoto, Makiko Seimiya, Malgorzata Santel, Barbara Treutlein, Franka Voigt

## Abstract

Spatiotemporal regulation of mRNA localisation is fundamental to cell identity specification and function, yet tracking transcript dynamics in living differentiating cells remains technically challenging. Here, we establish a robust pipeline for MS2 tagging of endogenous transcripts in human induced pluripotent stem cells (iPSCs), coupled with single-particle tracking and Hidden Markov Modelling to map mRNA mobility landscapes during differentiation and cell state transitions. Applying this approach to different cytoskeleton-encoding transcripts in diverse contexts — neural organoids, directly programmed neurons, and vascular organoids — we reveal a conserved principle: Both, *β*-actin and *β*2b-tubulin particle dynamics progressively shift towards constrained, compartmentalised patterns as cells acquire cell type identity. Perturbation experiments demonstrate that microtubule-dependent tethering is a common, conserved mechanism controlling *β*-actin mRNA localisation in all cell types studied, whereas translation-dependent anchoring and actin filaments contribute in a context-dependent manner. Analysis of particle dynamics in migrating blood vessel progenitors further showed that *β*-actin mRNAs accumulating at cell edges are highly diffusive, while those in perinuclear regions show constrained movement. Together, our integrated framework provides a scalable foundation for mechanistic dissection of mRNA targeting in human developmental and disease models.

## Main

Precise subcellular localisation of mRNAs enables cells to control where and when proteins are synthesised. This provides a spatiotemporal layer of gene expression regulation that is particularly critical during development, when cells must rapidly and asymmetrically reorganise their transcriptomes in response to instructive cues.^1–3^ Spatial targeting is especially important in highly polarised cells such as neurons, where local translation supports stimulus-specific responses at synapses far from the cell body.^4^ Systematic studies across diverse model systems - including Drosophila embryos, mammalian fibroblasts, intestinal epithelium, and neurons - have established that a substantial fraction of the transcriptome is non-uniformly distributed within cells, indicating that mRNA localisation is a pervasive feature of gene regulation rather than a specialised adaptation.^5–12^ Accordingly, dysregulation of mRNA localisation has been implicated in diseases such as neurodegeneration and cancer, underscoring its physiological importance.^13,14^ Many mRNA targeting mechanisms are conserved across polarized cell types.^12^ They often involve cis-regulatory sequences within mRNAs, termed “zipcodes”, which are recognised by trans-acting RNA-binding proteins (RBPs) that assemble transcripts into messenger ribonucleoprotein (mRNP) complexes. These complexes reach their destination via three broad mechanisms: directed transport along cytoskeletal networks; diffusion followed by entrapment at anchors; and selective protection from degradation. Targeting mechanisms are not mutually exclusive and often act synergistically, with mRNPs dynamically transitioning between active transport, passive diffusion, and stable anchoring to coordinate localisation with translational control.^15,16^ *β*-actin (ACTB) mRNA has become a paradigm for studying targeted mRNA localisation. A conserved 54-nt zipcode in its 3^*′*^ UTR is recognised by the RBP ZBP1 (human gene IGF2BP1), which is thought to mediate transport to sites of cytoskeletal remodeling.^17,18^ Although ZBP1 interacts with several kinesin family members,^19,20^ its precise role in transport remains uncertain: ZBP1 knock-down reduces *β*-actin mRNA recruitment and persistence at dendritic spines without impairing its long-range transport within dendritic segments, suggesting additional regulatory inputs.^21^ In vitro reconstitution has further shown that *β*-actin and *β*2b-tubulin (TUBB2B) mRNAs can be transported via an alternative pathway in which the RBP APC recruits transcripts to the kinesin adaptor KAP3, enabling processive transport over tens of micrometres.^22,23^ Consistent with a functional role for this mechanism, disruption of the APC–*β*2b-tubulin interaction impairs dynamic microtubule formation and cortical neuron migration in rat embryos.^24^ Together, these observations indicate that cytoskeletal mRNA localisation is not governed by a single RBP-RNA interaction, but requires the integration of multiple regulatory inputs whose relative contributions remain poorly defined, and whose importance in human cells has not been directly examined. Dynamic regulation of mRNA targeting is particularly important during growth and development, when differentiating cells continuously reorganise their cytoskeleton and local transcriptomes.^25^ Cells must dynamically redistribute mRNPs in response to developmental cues, and environmental stimuli,^26,27^ yet how mRNP mobility is modulated across human cell types and differentiation trajectories remains largely unexplored. Current approaches are constrained by reliance on non-human model systems, ectopic reporter constructs, and limited throughput that precludes comparative analysis across cell states. Crucially, no framework exists for tracking endogenous transcript mobility in living human cells across a developmental trajectory at single-molecule resolution. Here, we present a live single-molecule imaging strategy that resolves mRNA mobility across spatial, temporal, and developmental contexts in diverse human cell types. We focus on *β*-actin and *β*2b-tubulin as well-characterised, ubiquitously expressed transcripts subject to multi-layered localisation control, providing an informative entry point for dissecting targeting mechanisms in human cells. Using induced pluripotent stem cell (iPSC)-derived systems — including neural and vascular organoids as well as 2-dimensional cultures of polarised neurons — we establish a scalable live single-molecule imaging framework that resolves mRNA mobility across cell types and developmental stages within a defined human genetic background. This framework extends mechanistic insights from classical model organisms to human developmental biology, opening new avenues for understanding mRNA targeting in health and disease.

## Results

### Endogenous tagging of mRNAs for live single-molecule imaging in human iPSCs

To dissect mRNA targeting dynamics across human cell types and developmental stages, we established a live single-molecule imaging platform in the well characterised WTC-11 human iPSC line.^28^ Because of their ubiquitous expression, precisely defined localisation signals and central roles in cytoskeletal organisation, we focused on *β*-actin and *β*2b-tubulin as target transcripts. We employed the MS2–MCP system,^29^ in which stem-loop arrays inserted into the transcript of interest are bound by fluorescently tagged MS2 coat proteins (MCPs), enabling real-time visualisation of individual mRNA particles. To ensure imaging fidelity across differentiated cell types, all components were integrated endogenously, avoiding artefacts associated with overexpression of either the target transcript or the MCP reporter. To generate imaging cell lines, we first heterozygously integrated a Halo-tagged MS2 coat protein (MCP-Halo)^30^ into the AAVS1 safe harbour locus, chosen for its transcriptional permissiveness across cell types and its resistance to silencing upon differentiation.^31^ To facilitate flexible cassette exchange, the insert was flanked by Bxb1 recombination sites. Correct heterozygous integration was confirmed by polymerase chain reaction (PCR) across the integration junctions and by single-molecule fluorescent in-situ hybridization (smFISH) against the MCP-Halo transcript, which revealed a single transcription start site per cell (Figure S1a,b). MS2 stem loop arrays (see Methods) for visualization of target mRNAs were then inserted 30 bp downstream of the STOP codon in the 3’-UTR of either ACTB or TUBB2B using CRISPR/Cas9 gene editing.^28^ To enable selection of correctly edited cells, an EGFP expression cassette was introduced downstream of the stem loop array. The cassette was flanked by loxP recombination sites for Cre-mediated removal after selection (Figure 1a). Correct integration of the MS2 array was confirmed using PCRs spanning the integration junctions or the entire construct (Figure S1c), as well as Sanger sequencing of the insertion PCR product (Figure S1d). Given the high expression levels of ACTB and TUBB2B genes, we selected heterozygous clones (Figure S1b) to reduce labelled mRNA density during live imaging. We confirmed heterozygosity by dual-colour smFISH, which revealed a single MS2-positive transcription start site per cell (Figure S1e). Finally, *β*-actin-MS2 transcript integrity was verified by Northern blotting with probes against the ORF as well as the MS2 array, confirming the absence of alternative RNA species (Figure S1f). To establish an mRNA mobility analysis pipeline, we first performed live imaging in undifferentiated human iPSCs as a well-defined cellular reference state. Cells were labeled with the HaloTag ligand JF 549^32^ and image series were acquired at 20 Hz. As expected, individual mRNAs appeared as diffraction-limited spots with high signal-to-background in the cytoplasm, while unbound MCP-Halo protein accumulated in the nucleus (Figure 1b, Supplementary movie 1). To quantify the live single-molecule data, we developed a semi-automated image analysis pipeline for high-throughput particle tracking and diffusion coefficient extraction (see Methods). Across both cell lines, this approach yielded mobility features for 51,363 mRNA particles in total (representative tracks in Figure 1b, see Supplementary table 1), enabling quantitative analysis at a scale not previously accessible for live single-molecule mRNA data in human cells.

**Figure 1.**
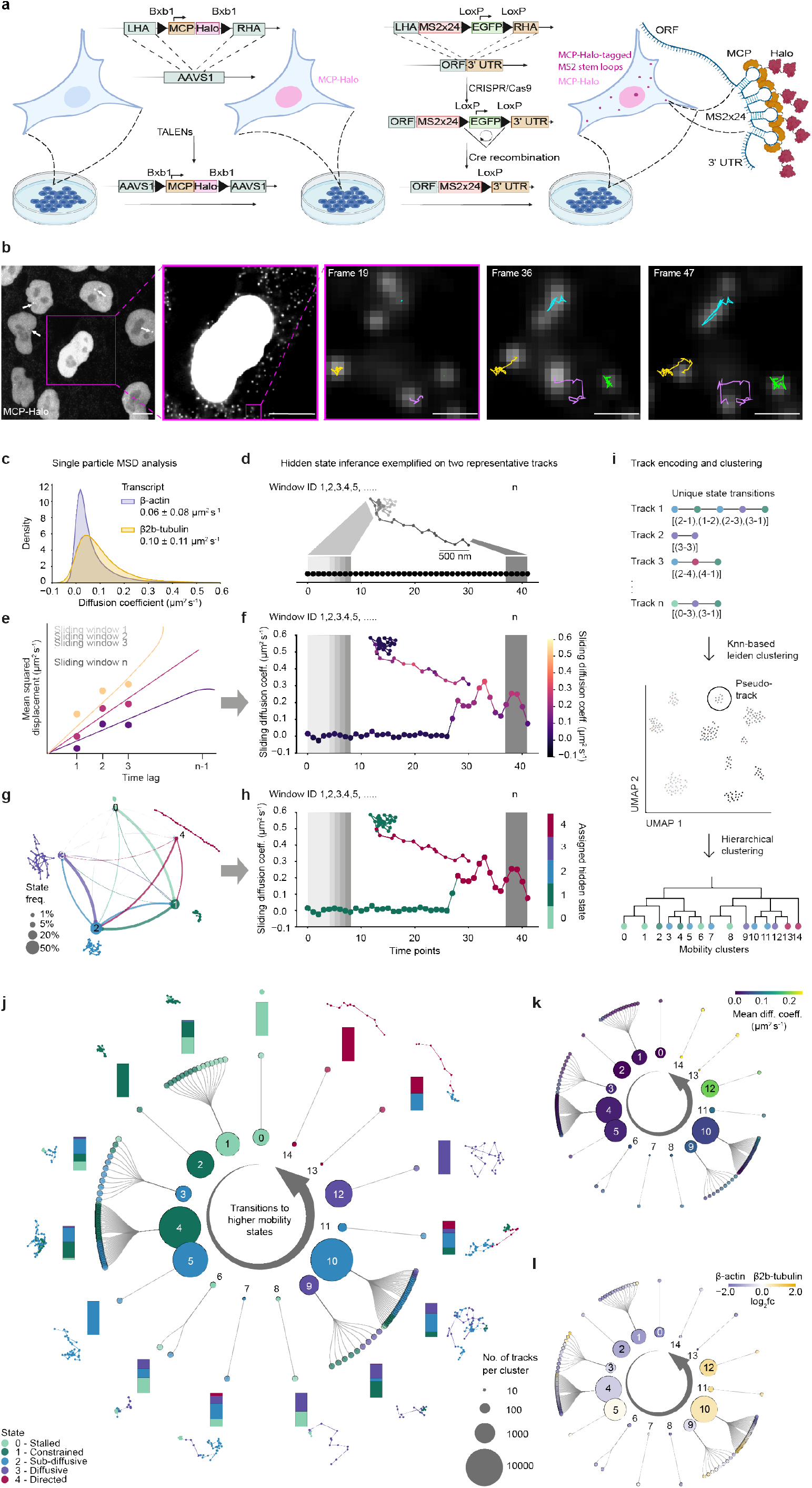
Live single-molecule imaging and Hidden Markov modelling of endogenously tagged transcripts uncover distinct mRNA mobility clusters in human iPSCs. **a**. Schematic of iPSC line generation. MCP-Halo was inserted into the AAVS1 safe-harbour locus using TALENs, and 24× MS2 stem loops were introduced into endogenous genes of interest by CRISPR/Cas9-mediated homology-directed repair. LHA, left homology arm; RHA, right homology arm. **b**. Single-molecule imaging of *β*-actin mRNA tagged with 24 MS2 stem loops. Left: Maximum-intensity projection of an iPSC colony. Nuclear accumulation of MCP-Halo due to an NLS and single transcription sites (white arrows) of the heterozygously tagged ACTB gene are visible. Inset 1: Zoom-in of a single cell (denoised image), where each diffraction-limited spot corresponds to one *β*-actin mRNA molecule. Inset 2: Selected frames from a 5 s live movie (20 Hz) overlaid with single-molecule tracks. **c**. Density plot of diffusion coefficients derived from the slope of single-track MSD curves. Diffusion coefficients are summarised as mean ± SD per transcript. **d**. Schematic of the sliding-window approach. Individual tracks were subdivided into overlapping sub-tracks using a window size of 5 consecutive time points and a step size of 1. Different shades of grey denote individual sliding windows. **e**. Extracting the diffusion coefficient per sub-track from the slope of a linear curve fitted to the first three time lags in the MSD plot. **f**. Resulting sliding diffusion coefficients for one representative track, where each diffusion coefficient was assigned to the coordinate pair in the centre of the window. **g**. Transition diagram showing the frequency of hidden states identified by the HMM (node size) and transition probabilities between states (arrow thickness). Example sub-tracks are shown for each state: 0: “stalled”, 1: “constrained”, 2: “sub-diffusive”, 3: “diffusive”, 4: “directed. **h**. States inferred by HMM from sliding diffusion coefficient vs. time plot of one representative track. **i**. Schematic of track encoding and clustering. Individual tracks were encoded by the relative frequency of unique state transitions and aggregated into pseudo-tracks using kNN-based Leiden clustering. Pseudo-tracks were further classified into top-level mobility categories by hierarchical clustering. **j**. Lineage-tree-like representation of cumulative *β*-actin and *β*2b-tubulin mobility classes in human iPSCs. Inner ring: Top-level clusters ordered by increasing di-grams (transitions to higher-mobility states) and coloured by the most frequent hidden state; node size indicates the number of tracks per cluster. Outer ring: Pseudo-tracks assigned to each mobility cluster, coloured by their most frequent hidden state. Stacked bars indicate the fraction of hidden states per cluster. Representative tracks are shown. **k**. Mean diffusion coefficient per cluster (inner circle) and pseudo-track (outer circle). **l**. Log_2_ fold-change in the number of *β*2b-tubulin tracks per cluster (inner ring) and pseudo-track (outer ring) relative to *β*-actin tracks.

### Hidden Markov Modelling of single-particle tracks reveals multiple transient mobility classes in human iPSCs

To characterise mRNA mobility in our dataset, we first calculated single-molecule diffusion coefficients (D) as a standard measure of molecular mobility, derived from mean squared displacement (MSD) analysis.^33^ Consistent with partially constrained cytoplasmic movement, we obtained D = 0.06 ± 0.08 *µ*m^2^s^-1^ and 0.10 ± 0.11 *µ*m^2^s^-1^ (mean ± SD) for *β*-actin and *β*2b-tubulin transcripts, respectively (Figure 1c). With standard deviations as large as the means themselves, these results suggested that a single average value would obscure substantial variability in the data. Indeed, closer inspection revealed two sources of heterogeneity: mRNA particles differed considerably from one another in their overall mobility, and — critically — individual particles did not move in a single consistent way, but instead switched between phases of constrained and more mobile behaviour over time. Averaging D over an entire trajectory would collapse these distinct behaviours into a single value obscuring biologically relevant information. Since particle mobility and mobility state switching can inform on the underlying molecular mechanisms that govern transcript localisation and processing, we developed a sub-track analysis approach that preserves this temporal detail. Rather than analyzing each trajectory as a whole, we used a sliding window to divide trajectories into short, overlapping segments and calculated D independently for each segment (Figure 1d, e). By mapping these local D values back onto the particle’s position and time, we generated time-resolved mobility profiles for individual mRNAs that reliably captured distinct mobility phases and the transitions between them within single trajectories (Figure 1f). To infer biologically relevant underlying mobility states from these time-resolved profiles, we used the sub-track D values as input for Hidden Markov Modelling (HMM), which has previously been applied to extract transient mobility states from noisy single-molecule data.^34^ Using a two-step hierarchical Gaussian HMM (see Methods), we identified four hidden states termed “stalled”, “constrained”, “sub-diffusive”, and “diffusive” together with their occurrence frequencies and transition probabilities (Figure 1g,h). Since periods of directed movement were not captured by the model, likely due to their rare occurrence, we retroactively introduced a fifth state (“directed”) comprising all sub-tracks that exhibited linear motion (*α* > 1.7 and net directionality > 0.85, see Methods). Leveraging these five mobility states, we next devised an approach to classify particles according to their mobility state sequences (see Methods). To this end, each track was encoded by its unique state transitions, weighted by their relative per-track frequencies (Figure 1i, top). The resulting feature vectors were used to construct a k-Nearest-Neighbour (kNN) graph that was partitioned into 175 clusters using the Leiden algorithm. These clusters, termed “pseudo-tracks”, each represented a unique combination of mobility state transitions and their frequencies (Figure 1i, middle). Pseudo-tracks with overlapping features were then aggregated into 15 mobility classes using hierarchical clustering (Figure 1i, bottom). The resulting particle mobility map summarizes, for each cluster, the frequency of each mobility state (Figure 1j, stacked bar plots), the predominant state at the cluster level (Figure 1j, inner ring) and the predominant state at the level of individual tracks (Figure 1j, outer ring). Together, our analysis resolves the population-level averages seen in Figure 1c into biologically interpretable movement patterns, illustrated by representative tracks colored by mobility state (Figure 1h). Applying this framework to *β*-actin and *β*2b-tubulin transcripts in undifferentiated human iPSCs, we found that the majority of tracks (70.7 %) displayed mixed mobility behaviours, while only approximately 30 % of tracks were composed of a single state (clusters 0, 2, 5, 12 and 14). Intriguingly, relatively few tracks exhibited either highly diffusive (6.1 %, cluster 12) or directed (clusters 9, 13 and 14) behaviour. This suggests that even in undifferentiated stem cells most mRNAs are subject to tethering and experience heterogeneous, confined microenvironments. To assess whether mobility patterns were transcript-specific, we next compared the state distributions of *β*-actin and *β*2b-tubulin mRNAs by calculating the log_2_ fold-change in their relative contributions per cluster in undifferentiated iPSCs (Figure 1l). *β*-actin transcripts were predominantly enriched in lower-mobility clusters, consistent with their tethering for transport to sites of local translation and cytoskeletal remodeling,^17,18^ and the established link between elevated actin dynamics and pluripotency.^35^ *β*2b-tubulin transcripts, by contrast, were enriched in sub-diffusive and diffusive clusters, consistent with the fact that tubulin isotype TUBB2B predominantly plays a role in microtubule function during neuronal differentiation.^36^ Together, these differences reveal transcript-specific modes of regulation in human iPSCs that were not apparent from population-level diffusion coefficient measurements and could only be resolved using the hierarchical HMM-based framework described above.

### Live single-mRNA imaging in human neural organoids reveals cell-type specific shifts in particle mobility across neuronal differentiation

Using the established workflow, we next determined how mRNA mobility patterns change upon stem cell differentiation. We differentiated human iPSCs into neural organoids — in vitro self-organizing tissues that recapitulate key features of early human brain development. They generate distinct cell types — including neuroepithelial progenitors, radial glia, and neurons^37^ across a defined differentiation time course (Figure 2a). To enable single-molecule imaging throughout these large, opaque specimens, organoids were live-sliced into 200 *µ*m thick sections (Figure 2b and c, see also Methods), a procedure previously shown to preserve tissue architecture while improving oxygen and nutrient diffusion.^38^ To control against detection of artifacts and establish a robust live single-molecule imaging pipeline, we generated organoids from the parental cell line that expresses only the MCP-Halo construct. At week 12, these specimens showed evenly distributed nuclear MCP-Halo fluorescence with no diffraction limited spots in the cytoplasm (Figure S2a,b), confirming that the cytoplasmic puncta observed in subsequent experiments are genuine mRNA particles. To delineate individual cell boundaries and capture cell-type specific mRNA dynamics at defined developmental stages, we generated mosaic organoids in which *β*-actin- or *β*2b-tubulin-MS2 tagged iPSCs were interspersed with wild-type (wt) cells (Figure 2a,d). To assess cell viability after slicing, we co-stained mosaic organoid sections with DRAQ7, a live-dead dye that only penetrates disintegrated nuclear membranes, and confirmed that tissue slicing did not compromise cell viability (Figure S2c). To verify that mosaic organoids developed normally and that tagged cells showed no differentiation bias, we performed single-cell RNA sequencing, which confirmed the presence of expected cell types and regional identities (Figure S2d). Uniform distribution of MCP-Halo-expressing cells across the UMAP confirmed that engineered cells were represented across all major cell populations (Figure S2e). Together, these results validate mosaic organoid slice cultures as a suitable system for live single-molecule imaging of mRNA dynamics during human neuronal differentiation. Thus, we next quantified *β*-actin and *β*2b-tubulin transcript mobility across three developmental time points - week 3, 5 and 8 (W3, W5, W8) - representing distinct stages of organoid development (Figure 2e, Supplementary movies 2-7, Supplementary table 1). While W3 organoids consist predominantly of radially-oriented progenitor cells, W5 organoids already contain immature neurons oriented perpendicular to the progenitors axis. By W8, the neuronal population has substantially expanded.^37^ Across this developmental trajectory, we observed a progressive global shift of particle mobility from highly diffusive towards more constrained states. At W3, when organoids were mainly comprised of progenitor cells, *β*-actin and *β*2b-tubulin particles were predominantly subdiffusive and diffusive. By W5, as the first immature neurons appeared, a shift towards more constrained mobility states became apparent. At W8, when the neuronal population had substantially expanded, particles were strongly enriched in stalled and constrained states (Figure S3b). To determine whether this shift reflected cell-type-specific regulation rather than a generic temporal effect, we assigned individual fields-of-view to progenitor (all time points), W5 neuron, or W8 neuron categories based on cell morphology and tissue architecture. We then computed mobility cluster enrichments for each cell type (Figure 2f). This analysis revealed a clear cell-type-specific pattern: Despite their established polarity axis, progenitors were strongly enriched in high-mobility populations (clusters 5, subdiffusive; 9 and 12, diffusive). W5 neurons occupied a transitional state, showing enrichment in clusters 6 and 7 that are characterised by switching between mobile and tethered phases, and suggesting that the shift towards constraint is an active and regulated process. W8 neurons were predominantly enriched for stalled or constrained particles (clusters 0 and 2). Interestingly, cluster 14 (directed motion particles) appeared enriched in progenitors and W5 neurons, though because of its small track count (< 100 tracks, see node sizes in Figures 2f and S3b) this observation warrants cautious interpretation. Unlike the differential regulation observed between *β*-actin and *β*2b-tubulin in iPSCs, the two transcripts showed broadly similar mobility profiles across all three neuronal cell types, suggesting convergence onto shared targeting mechanisms during neuronal differentiation. To assess whether apparently stalled particles might represent slow transport complexes moving below our detection threshold, we reduced the sampling frequency to 2 Hz and extended movies to 50 s. We find that state distributions were largely preserved across sampling rates (Figure S4a,b), confirming that cell-type-dependent mobility differences are robust (Figure S4c). Together, these results demonstrate that mRNA mobility is dynamically and cell-type-specifically regulated across human neuronal differentiation. While progenitors maintain high transcript diffusivity, immature neurons occupy a transitional mobility state, and mature neurons display tightly constrained transcript localisation, establishing a progressive developmental trajectory from mobility to constraint. The similarity between *β*-actin and *β*2b-tubulin mobility profiles across this trajectory suggests shared cytoskeletal targeting mechanisms become dominant as cells acquire neuronal identity.

**Figure 2.**
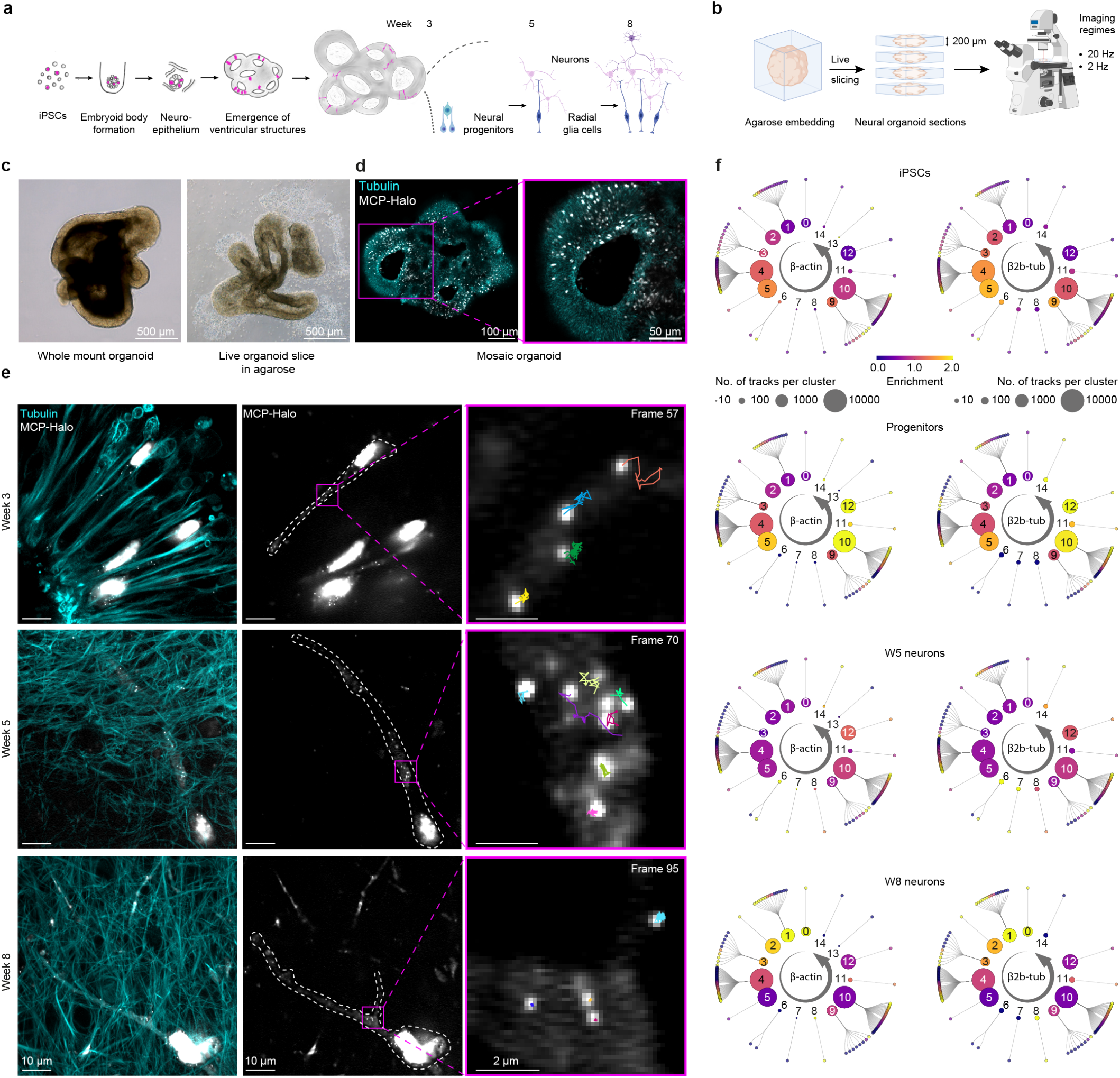
Live single mRNA imaging in neural organoids reveals global shifts in mobility cluster compositions throughout differentiation. **a**. Schematic of mosaic neural organoid generation. Magenta indicates cells genetically modified for live imaging, grey denotes unmodified cells. Transgenic cells were mixed with wt cells at a ratio of 5–15 %. **b**. Vibratome sectioning of live neural organoids embedded in low-melting agarose. Organoids were sectioned into 200 *µ*m slices, directly mounted on glass coverslips, and imaged at two different sampling speeds. **c**. Bright-field image of a three-week-old whole-mount organoid. Right: 200 *µ*m thick section of the same organoid. **d**. Left: Confocal image of a live mosaic organoid section. Right: Zoom-in of a single ventricle, showing MCP-Halo-labelled cells interspersed with wt cells; tissue architecture is outlined by a live tubulin dye. **e**. Live imaging of tissue sections at three developmental time points (denoised images). ACTB-MS2 labelled cells interspersed with wt cells are outlined in white. Insets show individual molecules and their corresponding tracks. **f**. Mobility cluster and pseudo-track enrichment for tagged *β*-actin (left) and *β*2b-tubulin (right) transcripts across different cell types. Enrichment was calculated as the fraction of particles in a given cluster or pseudo-track within one cell type divided by the average fraction across all cell types. A value of 1 indicates neither enrichment nor depletion.

### *β*-actin mRNA targeting patterns in human induced neurons

To corroborate our findings from organoid imaging and facilitate perturbation experiments, we next turned to human NGN2-induced neurons (iNeurons) as a complementary 2D system, focusing on *β*-actin because of its representative mobility profile. To this end, we generated ACTB-MS2 and wt iPSCs stably expressing a doxycycline-inducible NGN2 transgene (Figure 3a). NGN2 is a pioneer transcription factor that, upon overexpression, rapidly drives neuronal fate, largely bypassing a canonical progenitor stage with cells acquiring neuronal morphology as early as three days after induction.^39^ As in the organoid experiments, we induced mosaic cultures of ACTB-MS2 and wt cells. We confirmed MS2 and ACTB-ORF co-localisation via smFISH (Figure S5a) and verified neuronal identity using immunofluorescence (IF) staining (Figure 3b). As expected, all cells were MAP2-positive, and strong peripherin (PRPH) staining was consistent with previous reports of NGN2-induced cultures consisting predominantly of peripheral nervous system neurons.^40^ We performed live imaging at days 3, 5, and 10 post induction — time points chosen to approximate the organoid stages analyzed above. To enable mechanistic dissection of cytoskeletal contributions, we co-stained with live dyes for tubulin and filamentous actin and quantified *β*-actin mRNA association with individual cytoskeletal elements during differentiation-induced remodeling (Figure 3c, Supplementary movies 8-10, Supplementary table 1). *β*-actin mRNA mobility in iNeurons transiently increased at day 3 — reflected by the enrichment of diffusive clusters 10 and 12 — before progressively shifting towards more constrained behaviour by day 10 (Figure 3d). This recapitulates the early mobility burst and subsequent constraint observed across neuronal differentiation in organoids. Although the lowest-mobility clusters prominent in W8 organoid neurons were not yet apparent at day 10, we anticipate that continued maturation would reveal a further shift towards greater constraint. *β*-actin transcripts have been reported to localise to axonal growth cones, dendritic spines, and branch points for local translation.^21,41,42^ We therefore leveraged our mosaic cultures to ask whether the mobility patterns we identified were compartment-specific, automatically segmenting individual cells into soma, neurite, and outgrowth regions (Figure S5b). Consistent with the literature, particles from lower-mobility clusters (0, 1, 3 and 4) and the actively transported cluster (cluster 11) were predominantly localised in neurites, consistent with tethering to cytoskeletal elements for transport to and translation at distal sites (Figure S5c). Closer inspection revealed additional layers of spatiotemporal regulation. Fully stalled particles (cluster 0) showed a striking compartment bias that shifted over time: at day 3 they were largely confined to neurites, while by day 10 they were evenly distributed between neurites and soma. Highly diffusive particles (cluster 12) showed the opposite trend — enriched in soma and outgrowths at day 3, they progressively redistributed toward neurites and outgrowths by day 10. Together, these compartment- and time point-specific patterns suggest that *β*-actin mRNA dynamics are governed by multiple targeting mechanisms operating in parallel across distinct cellular compartments, and that the balance between these mechanisms shifts as neurons mature. The heterogeneity in these targeting patterns may reflect the diversity of ribonucleoprotein complexes that *β*-actin mRNA assembles into. While long-range transport of *β*-actin Mrna is thought to be predominantly kinesin- and microtubule-dependent,^19,20,22,23^ short-range positioning and anchoring in growth cones, axons and dendritic spines is proposed to rely on myosin motors and actin filaments.^21,43–45^ To quantify the relative contributions of these cytoskeletal elements to *β*-actin mRNA tethering, we measured F-actin and microtubule fluorescent dye intensities in pixel positions occupied by *β*-actin mRNA particles during live imaging (Figure S5d, top). Interestingly, we found that stalled and low-mobility particles (cluster 0 and 1-3) strongly co-localised with both microtubules and actin filaments, while particles from the directed-transport cluster 11 preferentially co-localised with actin filaments.

**Figure 3.**
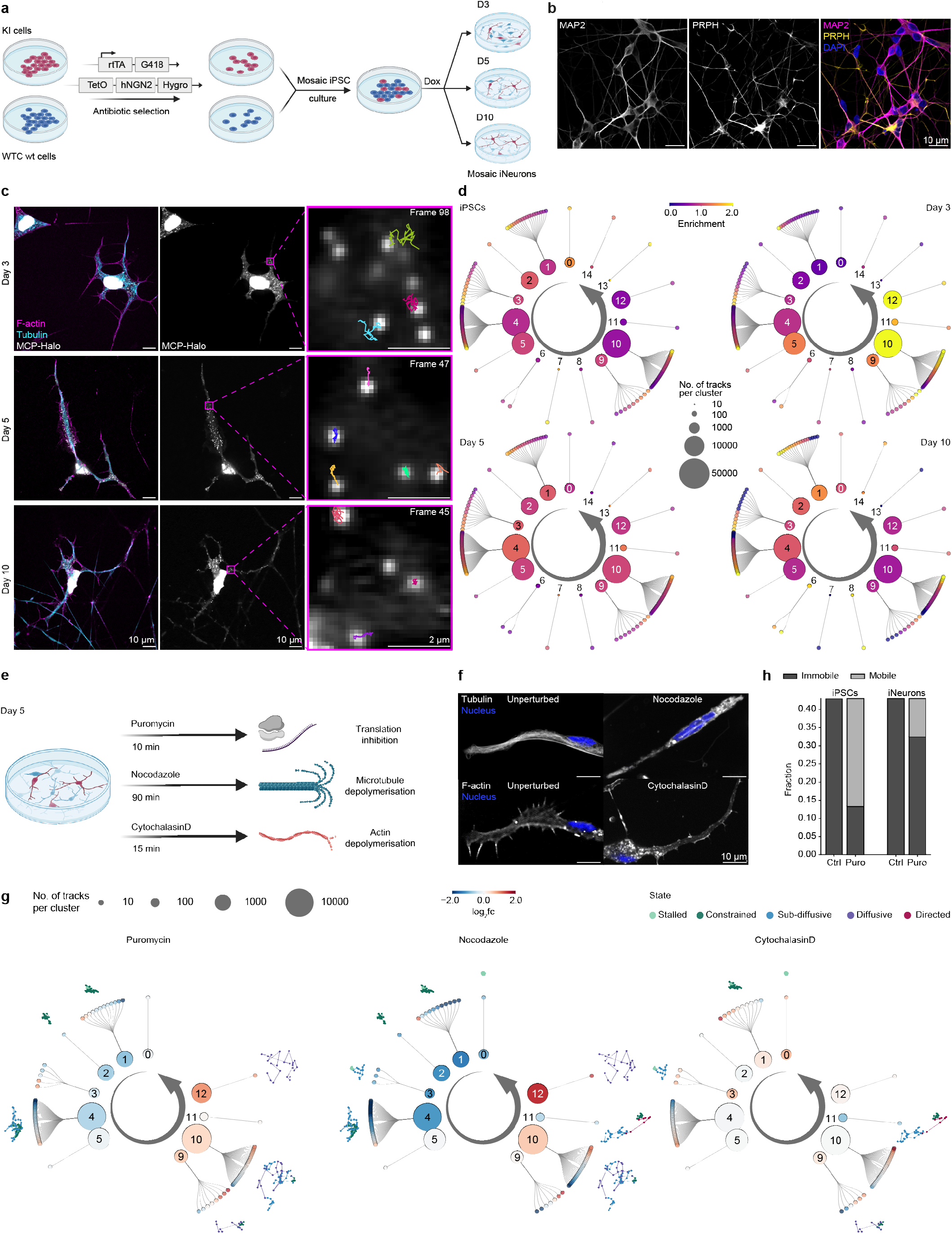
Live single mRNA imaging in iNeurons recapitulates global mobility shifts and identifies microtubules as key regulators of *β*-actin targeting during neuronal differentiation. **a**. Schematic of iNeuron generation and experimental design. Transgenic and wt cells were co-infected with two viruses encoding a constitutively active rtTA and doxycycline-inducible human NGN2, respectively. Cells were selected via antibiotic resistance, single colonies were isolated, and iNeurons were generated from a mosaic culture consisting of 30 % modified transgenic and 70 % modified wt cells. **b**. Antibody staining of iNeuron cultures at day 10: MAP2 marks neurons and is enriched in dendrites, PRPH marks neurons of the peripheral nervous system. **c**. Live imaging of mosaic iNeuron cultures at three developmental time points (denoised images). The cytoskeleton is labelled with live tubulin and F-actin dyes (left panel), with selected single-molecule tracks shown in the inset (right panel). **d**. Quantification of mobility cluster and pseudo-track enrichment for tagged *β*-actin transcripts across developmental time points. Enrichment was calculated as the fraction of particles in a given cluster or pseudo-track at one time point divided by the average fraction across all time points. **e**. Schematic of small-molecule perturbation experiments. Day 5 iNeurons were treated with small molecules and imaged after the indicated treatment durations. **f**. Top: Live tubulin staining in untreated cells (left) and cells treated with 10 *µ*M nocodazole (right). Bottom: Live F-actin staining in untreated cells (left) and cells treated with 10 *µ*M cytochalasin D (right). For nocodazole and cytochalasin D, incubation times were optimised to achieve efficient depolymerisation of the respective components (as seen by diffuse live dye signal), while still keeping the overall cell shape mostly intact. **g**. Top: Log fold-change showing particle fraction enrichment/depletion per cluster and pseudo-track in perturbed vs. unperturbed conditions. Bottom: Representative tracks from the most enriched and depleted clusters. **h**. Fraction of immobile vs. mobile particles (aggregation of states 0 and 1, and states 2 and 3, respectively) in unperturbed and puromycin-treated conditions in iPSCs and day 5 iNeurons.

### Translation-dependent tethering and actin filaments play a secondary role in *β*-actin mRNA targeting during neuronal differentiation

Tethering of mRNAs at specific subcellular sites is thought to be controlled by a combination of cytoskeletal anchoring and ribosome engagement at sites of active translation.^46^ To dissect the relative contributions of cytoskeletal structure and active translation to the tethering of *β*-actin mRNAs in iNeurons, we performed a series of small molecule perturbation experiments. We treated cells with either puromycin (Puro, translation inhibition^47,48^), nocodazole (Noco, microtubule depolymerisation^49^) or cytochalasin D (CytoD, actin depolymerisation^50^) (Figure 3e,f, Supplementary movies 11-13) and computed cluster enrichment as log_2_ fold-change relative to the unperturbed condition (Figure 3g). Puro treatment increased the fraction of particles in the most diffusive clusters (clusters 9, 10, and 12; log_2_fc = 0.48, 0.39, and 0.87 respectively), confirming translation-dependent anchoring of a subset of *β*-actin mRNA molecules. However, fully stalled particles (cluster 0) were preferentially associated with tubulin (Figure S5d, middle) and largely unaffected by Puro treatment, suggesting that these transcripts are not part of locally tethered actively-translating nascent chain complexes. Notably, the overall effect of Puro was modest: partitioning of particles into immobile and mobile fractions revealed only a small increase in mobility upon treatment (55.3 % to 67.5 %; Figure S5e). This stands in marked contrast to iPSCs, where Puro treatment produced a far more pronounced increase in the mobile fraction (56.4 % to 86.7 %; Figure 3h), and suggests that ribosome-mediated tethering plays a substantially reduced role in maturing neurons compared to pluripotent cells. Microtubule depolymerisation with Noco produced considerably larger effects, strongly depleting low-mobility clusters including cluster 0 (log_2_fc = 0.94) and enriching diffusive clusters 10 and 12 (log_2_fc = 0.50 and 1.47 respectively) (Figure 3g), with a corresponding increase in the mobile fraction from 55.3 % to 77.0 % (Figure S5e). The directed-transport population (cluster 11) was also depleted (log_2_fc = 0.54), consistent with microtubule-dependent long-range transport of *β*-actin mRNA.^19,20,22,23^ Actin depolymerisation with CytoD produced the most unexpected result: rather than releasing tethered particles, loss of the actin cytoskeleton increased the abundance of several low-mobility clusters (clusters 0 and 3; log_2_fc = 0.64 and 0.55, respectively), while the overall change in the immobile fraction was negligible (44.7 % to 46.0 %; Figure S5e). Together, these data implicate microtubule anchoring as the dominant mechanism of *β*-actin mRNA tethering in differentiating neurons. In summary, these perturbation experiments demonstrate that *β*-actin mRNA targeting in iNeurons is dynamically regulated along the differentiation trajectory and exhibits compartment-specific features. Multiple pathways act in concert to regulate *β*-actin mRNA localisation, collectively fine-tuning both the precise subcellular location and translation status of individual transcripts. We suggest that active translation and interactions with the actin cytoskeleton play secondary roles in this process, with microtubule-dependent anchoring emerging as the dominant targeting mechanism during neuronal differentiation.

### Imaging *β*-actin mRNA dynamics in sprouting human blood vessel progenitors

Besides neurons, polarised migratory cells are another widely used model for studying mRNA targeting. Indeed, fundamental targeting mechanisms have been proposed to be conserved across cell types,^51^ raising the question of whether the microtubule-dominant pattern we observed in neurons extends to other polarised human cells. To further explore this we next assayed *β*-actin transcript dynamics in highly migratory blood vessel progenitor cells derived from ACTB-MS2 iPSCs. To this end, we generated sprouting blood vessel organoids (BVOs).^52,53^ BVOs consist of highly motile endothelial and mural cells migrating rapidly away from the colony centre and thus represent an ideal model system to study localised gene expression dynamics in polarised, migratory cells (Figure 4a). As before, we first used IF to confirm that our tagged cell line differentiated into the expected cell types, as indicated by mutually exclusive expression of PDGFR-*β* (mural cells) and CD31 (endothelial cells) (Figure 4b). We then acquired live imaging data to assess *β*-actin mRNA mobility at distinct time points (day 7 and 10) across the differentiation trajectory (Figure 4a,c, Supplementary movies 14,15, Supplementary table 1). Although both BVOs and neural model systems showed similar overall trends, changes in particle mobility across time points were much less pronounced in BVOs (Figure 4d). Early differentiation (day 7) showed only moderate enrichment of mobile clusters (clusters 10–12; enrichment score ∼1.3) compared to the stronger enrichment seen in day 3 iNeurons (∼1.5 to 2.0). Similarly, enrichment of constrained and sub-diffusive clusters (clusters 1–5) at day 10 was modest (enrichment score ∼1.1–1.3). Most strikingly, fully stalled particles (cluster 0) were significantly depleted at day 10 in BVOs, likely reflecting the highly dynamic and transient cytoskeletal architecture of these rapidly migrating cells. This contrasts with neuronal models, in which these fully stalled particles (cluster 0) frequently co-localised with cytoskeletal elements in a translation-independent manner. *β*-actin mRNA has previously been reported to accumulate at the leading edges of migratory cells, where local translation is thought to support actin filament assembly at sites of active cytoskeletal remodeling.^17,54^ To test whether this was reflected in cytoskeletal co-localisation patterns, we quantified the association of specific mobility clusters with individual cytoskeletal components (Figure 4e). Interestingly, co-localisation with actin filaments was rarely observed. Instead, particles in low-mobility clusters (clusters 0–4 and 6) showed strong association with the tubulin cytoskeleton. This points to a dominant role for microtubules in localizing *β*-actin mRNA during cell migration, consistent with our findings in neurons and suggesting that microtubule-dependent targeting may represent a conserved mechanism for *β*-actin targeting across polarised human cell types.

**Figure 4.**
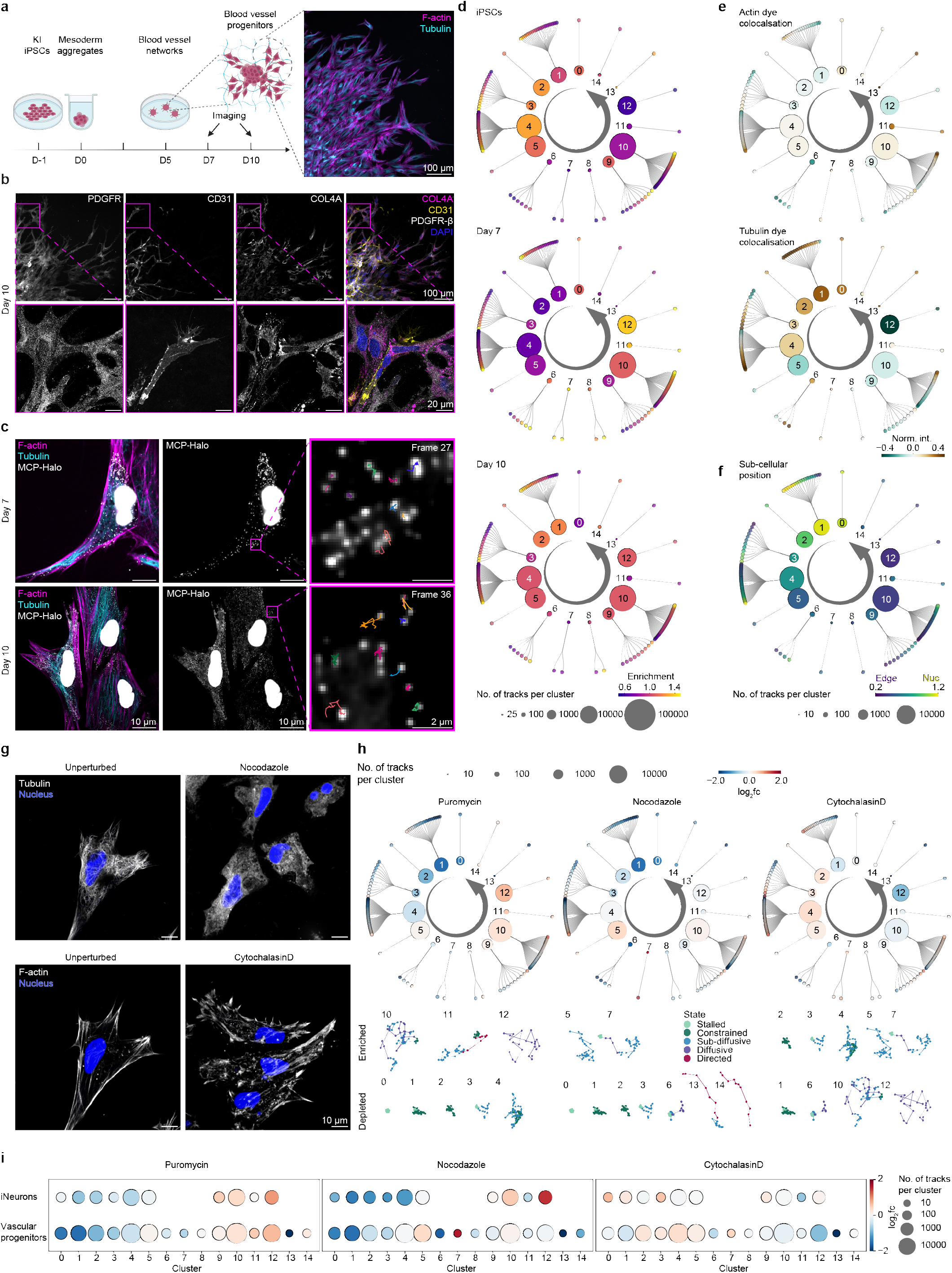
Tracking endogenously tagged *β*-actin mRNAs in human vascular organoids reveals subcellular localisation-dependent dynamics. **a**. Schematic of blood vessel progenitor generation. Left: Cells were aggregated and directed towards the mesoderm lineage, then plated into a solidifying Collagen I-Matrigel solution on day 5, in which vessel-like structures formed. Cells were stained and imaged directly within the hydrogel. Right: Confocal image of the edge of a blood vessel progenitor colony at day 10. Sprouting cells at the colony edge were selected for live imaging. **b**. Antibody staining of blood vessel progenitors at day 10: Endothelial cell network (CD31, yellow) covered by mural cells (PDGFR-*β*, white), embedded in a basement membrane (COL4A, magenta). **c**. Live imaging of sprouting blood vessel progenitors at two developmental time points, (100 X magnification, denoised images). The cytoskeleton is labelled with live tubulin and F-actin dyes (left panel), with selected single-molecule tracks shown in the inset (right panel). **d**. Quantification of mobility cluster and pseudo-track enrichment for tagged *β*-actin transcripts across developmental time points. Enrichment was calculated as the fraction of particles in a given cluster or pseudo-track at one time point divided by the average fraction across all time points. **e**. Z-score–normalised cytoskeletal intensities per track, averaged by cluster and pseudo-track in day 10 sprouting cells. For each detected spot ((x,y) coordinate pair), intensities from live-dye channels were extracted, normalised, and averaged per track, providing a proxy for mRNA co-localisation with the actin or tubulin cytoskeleton. **f**. Distance-to-edge/nucleus ratio, averaged by cluster or pseudo-track in day 10 sprouting cells. For each track, the distance to the cell edge (cell mask) divided by the distance to the nucleus (nucleus mask) was calculated. Higher values indicate proximity to the nucleus, lower values proximity to the cell edge. **g**. Top: Live tubulin staining in untreated cells (left) and cells treated with 10 *µ*M nocodazole for 45 min (right). Bottom: Live F-actin staining in untreated cells (left) and cells treated with 10 *µ*M cytochalasin D for 45 min (right). Diffusive live dye signal confirms depolymerisation of the respective cytoskeletal components. **h**. Top: _2_ fold-change per cluster or pseudo-track in perturbed versus unperturbed conditions. Bottom: Representative tracks from the most enriched and depleted clusters. **i**. Comparison of cluster enrichment and depletion between iNeurons and vascular progenitor cells across all perturbations.

### *β*-actin mRNAs are highly diffusive in the periphery of migratory cells

To further explore the microtubule-dominant localisation of *β*-actin mRNA in migratory cells, we quantified particle mobility with respect to subcellular localisation. Specifically, we computed the ratio of a particle’s distance from the cell edge to its distance from the nucleus. Surprisingly, high particle mobility strongly correlated with proximity to the cell edge (Figure 4f, dark color), while constrained particles localised closer to the nucleus (Figure 4f, bright color). Two explanations could account for this observation: First, *β*-actin mRNA transcripts may not get recruited to actin filaments for local translation at the cell periphery — if they did, a reduction in mobility at the cell edge would be observed. Second, the periphery may represent a biophysically distinct environment that imposes fewer constraints on particle movement. To quantify the impact of translation on *β*-actin mRNA mobility and further dissect the contributions of underlying tethering mechanisms, we again performed small molecule perturbations (Puro, Noco, CytoD; Supplementary movies 16-18) and assessed cytoskeleton depolymerisation sufficiency based on diffusive live dye signal (Figure 4g). As in iNeurons, translation inhibition by Puro treatment only moderately enriched high mobility clusters (10, 12) and slightly depleted low-mobility clusters (1-4) (Figure 4h,i). Particles remaining in low-mobility clusters localised further away from the nucleus (Figure S6a, left) and were anchored to both tubulin and actincytoskeleton (Figure S6b, left). This suggests that ribosome-dependent anchoring occurs closer to the cell centre, possibly at the endoplasmic reticulum. Globally, translation inhibition with Puro broadly mirrored the response observed in other cell types, increasing the mobile fraction (57.4 % unperturbed vs. 72.4 % Puro) and depleting low-mobility states (Figure S6c). Additional changes in directed and stalled populations suggest context-dependent coupling of translation status, cytoskeletal association, and subcellular positioning. Microtubule depolymerisation with Noco reinforced the dominant role of microtubules in *β*-actin mRNA tethering. It depleted low-mobility and directed clusters (Figure 4h,i) and shifted particles toward higher mobility states (57.4 % mobile in unperturbed vs. 69.3 % in Noco; Figure S6c). Specifically, low-mobility clusters 0 and 1 were strongly depleted (log_2_fc = 1.44 and 1.52), while the most mobile clusters (10 and 12) were largely unaffected (log_2_fc = 0.09 and 0.07, respectively). Directed-transport clusters (11, 13, and 14) were also depleted (log_2_fc = 0.22, 17.04, and 1.00; Figure 4h, i). Notably, the particles that remained anchored after nocodazole treatment were positioned closer to the nucleus than in the control condition (Figure S6a), suggesting that microtubule-dependent tethering preferentially occurs at the cell periphery. As in iNeurons, actin depolymerisation with CytoD had the least impact of all three perturbations and cumulative population mobility was essentially unchanged (57.1 % mobile in both unperturbed and CytoD; Figure S6c). Although the specific clusters affected differed between cell types (Figure 4h,i), residual low-mobility particles continued to co-localise with microtubules in both cell models (Figures S5d and S6b), further underscoring the primacy of microtubule-based tethering. One notable difference from iNeurons was the depletion of the most diffusive cluster (cluster 12; log_2_fc = 0.85) in vascular progenitor cells. Since highly mobile particles were preferentially located near the cell periphery, this depletion may reflect disruption of the high-mobility microenvironment normally maintained by the actin cortex at the leading edge. Upon CytoD treatment, particles may therefore have been redistributed into more microtubule- and ribosome-dense central regions. In summary, our data demonstrate a dominant role for microtubule-dependent targeting of *β*-actin mRNA across diverse iPSC-derived model systems. While cytoskeleton anchoring mechanisms appear conserved across cell types, the degree of co-translational anchoring differs in distinct differentiation states and cell types. Notably, highly mobile particles were enriched at the cell periphery and excluded from central regions in migratory cells. We hypothesise that these transcripts are targeted to the periphery to support dynamic actin cytoskeleton remodeling. How these transcripts remain enriched in proximity to peripheral actin networks, despite their high mobility, remains to be determined.

## Discussion

Live single-molecule imaging of endogenous transcripts in human stem cells has remained technically challenging, limiting mechanistic insight into how mRNA localisation is regulated during differentiation. By establishing a scalable MS2-based imaging platform in human iPSCs and coupling it with hierarchical Hidden Markov Modelling, we show that many *β*-actin and *β*2b-tubulin mRNA particles dynamically transition between distinct mobility states, rather than exhibiting a uniform diffusion rate, a level of behavioural complexity that ensemble-level approaches would obscure. Comparing *β*-actin and *β*2b-tubulin mRNA dynamics in iPSCs revealed that the two transcripts are subject to distinct modes of regulation at the pluripotent stage. While *β*-actin transcripts were predominantly constrained or tethered, *β*2b-tubulin particles were largely diffusive. This is consistent with the previously established, constraining function of ZBP1, which is highly expressed in iPSCs, and with the low, stochastic expression of APC, a key trans-acting regulator of *β*2b-tubulin localisation.^37^ Transcript-specific differences were no longer apparent during neuronal differentiation, where both *β*-actin and *β*2b-tubulin mRNAs showed similar mobility profiles, suggesting convergence towards shared targeting mechanisms as cells adopt a terminal fate. In neural progenitors both transcripts underwent a transient burst in mobility before progressively shifting towards constrained and stalled states in developing neurons. This pattern was reproducible across two imaging timescales and validated in both the neural organoid and iNeuron models. Perturbation experiments in induced neurons revealed that this progressive constraint is predominantly microtubule-dependent. Puromycin-mediated ribosome release had only a modest effect on particle mobility in iNeurons, in contrast to its pronounced effect in iPSCs. This finding is consistent with previous reports suggesting that mRNAs in neuronal granules are often stored in stalled polysomes that are not susceptible to Puro treatment.^56^ Our data illustrate the dynamic reorganisation of neuronal granules upon differentiation and indicate that *β*-actin mRNAs are not predominantly tethered as actively translating mRNPs in maturing neurons. Nocodazole treatment substantially increased the mobile fraction and depleted directed-component clusters, consistent with microtubule-dependent long-range transport and tethering of *β*-actin mRNA.^19,20,22,23^ In contrast, actin depolymerisation had comparatively little effect and residual low-mobility particles continued to associate with microtubules in both neuronal and vascular models. This suggests microtubule anchoring as a dominant and conserved targeting mechanism across the cell types and subcellular localisations examined. Notably, none of the perturbations fully mobilised the particle population, indicating that multiple mechanisms, both active and passive, likely operate in parallel to regulate *β*-actin mRNA tethering. Since *β*-actin mRNAs co-localise with F-actin at axonal growth cones, branch points and dendritic spines,^45,43,57,21^ dissecting mobility state as a function of sub-cellular compartment in more mature neuronal models remains an important research direction. In migratory vascular progenitor cells, *β*-actin mRNAs were again predominantly anchored on microtubules, not the actin cytoskeleton, and in a translation-dependent manner. However, overall particle dynamics differed markedly from neuronal systems: mobility changes across differentiation were more modest, and fully stalled particles were depleted at later stages, likely reflecting the highly dynamic cytoskeletal architecture of rapidly migrating cells. The most mobile particles were preferentially enriched at the cell periphery, consistent with their proposed recruitment to leading edges to support local actin cytoskeleton synthesis.^58,59^ Yet, the high mobility of these peripheral particles argues against stable anchoring for local translation, and instead suggests restriction by a structural element or biophysical property of the local cytoplasm. Together with the depletion of peripheral particles upon actin depolymerisation,^60^ these findings raise the possibility that the actin-rich cell periphery generates a permissive microenvironment that promotes transcript accumulation through diffusion barriers rather than stable molecular anchoring. How transcripts are locally retained despite their high mobility remains an important open question. A longstanding bottleneck in the field has been the absence of scalable, human cell-compatible systems for high-resolution live mRNA imaging across diverse biological contexts. By integrating endogenous CRISPR tagging with stable MCP-Halo expression from the AAVS1 safe harbour locus, high-throughput image processing, and HMM-based sub-track mobility analysis, this framework enables systematic classification of over 10^5^ particles (Supplementary table 1) into biologically interpretable mobility classes - a scale and resolution not previously achieved in human stem cell models. We anticipate this approach will be broadly applicable to mechanistic dissections of mRNA targeting in human developmental and disease contexts, including the study of dysregulated transcript localisation in neurodegeneration.

## Methods

### Cell and organoid culture

#### iPSC maintenance

WTC-11 iPSCs (wild-type, male; Coriell Institute for Medical Research: GM25256) were used as parental cell line for all generated cell lines. All cell lines were cultured in mTeSR+ (STEMCELL Technologies, 05825) supplemented with penicillin–streptomycin (P/S, 1:200, Gibco 15140122) on matrigel-coated plates (Corning 354277). Cells were split every 4-5 days with TrypLE Express (Gibco, 12605010) and media was supplemented with CEPT^61^ (1:1000, final concentrations as in the original study: chroman-1 (50 nM, MedChem Express HY-15392), trans-isrib (0.7 *µ*M, Tocris 5284), emricasan (5 *µ*M, Selleckchem S7775), poly peptides (1:1000, Sigma-Aldrich P8483)) for one day after splitting.

##### Neural organoids

All organoids were generated as follows (adapted from an unguided whole-brain organoid differentiation protocol^62^): 500 cells were seeded per well of an ultra low-attachment 96-well plate (Corning CLS7007) in mTeSR+ (with 1:1000 CEPT and 1:100 P/S) to generate embryoid bodies (EBs). Media was exchanged to fresh mTeSR+ with 1:100 P/S on day 2. On day 4, neural induction media with 2 % dissolved matrigel (Corning, 356231) was supplied and exchanged every other day. Organoids were transferred to 6-well plates (Corning, 3471) on day 11 and differentiation media -VitA with 1 % matrigel was supplied and media was changed every 3 days. On day 28, differentiation media +VitA was supplied and organoids were transferred to 6 cm dishes (TPP, 93060) and moved to the shaker (65 rpm). Media was changed every 3-4 days. From day 42 onwards, 1:200 matrigel was supplied to the media.

#### iNeurons

iNeuron culture was adapted from Frega et al.^39^ as follows: For live imaging, 30’000 cells (ibidi 4-well dish, 80427) or 15’000 cells (ibidi 8-well dish 80807) were seeded per well of a matrigel-coated plate in mTeSR+ with 1:100 P/S, 1:1000 CEPT and 4 *µ*g/ml doxycycline (Sigma-Aldrich D9891-5G). On day 1, media was changed to neural induction media, and on day 3 media was changed to differentiation media. Doxycycline was removed after day 3 and media was changed every other day. 1 *µ*M cytosine-D-arabinofuranoside (Ara-C, Sigma-Aldrich C1768-100MG) was added on day 5 in case undifferentiated /off-target cells started overgrowing the neurons.

#### Vascular progenitor cells

Blood vessel progenitor cells were generated as follows (adapted from Wimmer et al.^52^): Cells were seeded into Corning Elplasia round-bottom, ultra-low attachment plates (Corning, 4442) at a concentration 300 cells per microcavity in aggregation media. On day 0, media was exchanged to N2B27 media, which was supplemented with 2 *µ*M Forskolin (Sigma Aldrich, F3917) and 100 ng/ml VEGF-A (Peprotech, 10020) on day 3. On day 5, aggregates were embedded into Matrigel/collagen-l solution, pH 7.4, onto a glass-bottom 4-well (ibidi, 80427) or 8-well (ibidi, 80807) dish. From day 5, blood vessel networks were cultured in BVO media and media was exchanged every 2-3 days.

#### Media

##### Neural organoids

Neural induction media: DMEM/F12 (Gibco, 31330-038), 1:100 N2 supplement (Gibco, 17502-048), 1:100 Glutamax supplement (Gibco, 35050061), 1:100 MEM-NEAA (Sigma, M7145), 1:5000 heparin solution (5 mg ml^1^, Gibco, H3149-10KU), 1:100 P/S Differentiation media: 1:1 - DMEM/F12 : neurobasal (Gibco, 21103-049), 1:200 N2 supplement, 1:100 B27 with (Gibco, 17504-044) or without (Gibco, 12587-010) VitA supplement, 1:4000 insulin (Sigma, I9278), 55 *µ*M 2-mercaptoethanol solution (Gibco, 21985023),1:100 Glutamax supplement, 1:200 MEM-NEAA, 1:100 P/S

##### iNeurons

Neural induction media: DMEM/F-12 supplemented with 4 *µ*g/ml doxycycline (Sigma-Aldrich D5207-1G), 1:100 N-2 supplement, 1:100 MEM-NEAA, 10 ng/ml human BDNF (Pepro-Tech, 450–02), 10 ng/ml human NT-3 (PeproTech, 450-03) Differentiation media: Neurobasal medium with 1:50 B-27 supplement, 1:100 GlutaMAX supplement, 10 ng/ml human BDNF, 10 ng/ml human NT-3.

##### Blood vessel networks

Aggregation media: KnockOut DMEM/F12 (Gibco, 12660012), 20 % KnockOut Serum Replacement (Gibco, 10828028), 1 % GlutaMAX (Gibco, 35050061), 1 % NEAA (Gibco, 11140035), 55 *µ*M *β*-mercaptoethanol (Gibco, 21985023), 100 U/ml penicillin-streptomycin, P/S (Gibco, 15140122), 50 *µ*m Y-27632 (Stem Cell Technologies, 72304) N2B27 media: 1:1 - Neurobasal (Gibco, 21103049) : DMEM/F12 (Gibco, 11330031), 1:100 B27 supplement (Gibco, 12587010), 1:200 N2 supplement (Gibco, 17502048), 1:200 GlutaMAX (Gibco, 135050061), 55 *µ*M *β*-mercaptoethanol (Gibco, 21985023), 100 U/ml P/S (Gibco, 15140122), supplemented with 12 *µ*M CHIR99021 (Tocris Bioscience, 4423) and 30 ng/ml BMP4 (Miltenyi Biotech, 130111165) Collagen I-Matrigel solution: 11.3 % v/v 0.1 N NaOH (Sigma Aldrich, S2770), 4.7 % v/v 10X DMEM (prepared prior, Sigma Aldrich, D5648-10L), 0.9 % v/v HEPES buffer (Gibco, 15630080), 0.7 % v/v NaHCO3 (Gibco, 5080094), 0.5 % v/v GlutaMAX (Gibco, 25050061), 6.9 % v/v Ham’s F12 (Gibco, 11765054), 50 % v/v PureCol (Advanced BioMatrix,5005), 25 % v/v GFR Matrigel (Corning, 356231) BVO (blood vessel organoid) media: StemPro-34 SFM media, supplemented with StemPro-34 nutrient supplement, 1 % GlutaMAX, 100 U/ml penicillin-streptomycin, 15 % fetal bovine serum (FBS) (Sigma-Aldrich, F2442-500ML), 100 ng/ml VEGF-A (Peprotech, 10020) and 100 ng/ml FGF2 (Miltenyi Biotech, 1300930841)

### Cell line generation

#### AAVS1-MCP-Halo

The WTC AAVS1-MCP-Halo parental cell line, hereafter referred to as “AAVS1-MCP-Halo”, was generated by integration of the targeting vector (TV) into the AAVS1 locus with TALENs as follows: A total of 20 *µ*g of DNA were prepared, with 90 % TV and 5 % of each nuclease plasmid (Addgene 59025^63^ and 59026^63^). WTC wt cells were dissociated with TrypLE for 3 min at 37°C and counted with the Countess II (Life Technologies). 800 000 cells were nucleofected in 100 *µ*l reaction volume (including the DNA mix) with the P3 Primary Cell 4D-NucleofectorTM X Kit (Lonza V4XP-3024), using the “hES-H9” programme. Cells were seeded into 10 cm dishes with 1:1000 CEPT. On day 3 selection with puromycin (1 *µ*g/ml, InvivoGen, ant-pr-1) was started, media was changed every day and CEPT was kept on for 3 days. When clear colonies started emerging, CEPT was removed and colonies were picked 9 days after nucleofection. Single clones were grown up and DNA was extracted from a fraction of the cells with the DNeasy Blood and Tissue Kit (Qiagen, 69504). Genotyping by PCR was performed with the following primers: left integration junction: AAVS1-MCP-LHA_fwd2 and AAVS1-MCP-LHA_rev; right integration junction: AAVS1-MCP-RHA_fwd and AAVS1-MCP-RHA_rev2; insert/wt band: AAVS1-MCP-LHA_fwd2 and AAVS1-MCP-RHA_rev2. PCRs were conducted with the Phusion™ High-Fidelity PCR Master Mix with HF Buffer (NEB, M0531S) according to the manufacturer’s protocol (https://www.neb.com/en/protocols/pcr-protocol-m0530?srsltid=AfmBOorXyKxT6q-Lj-V34jj44p3vIXX1QLATQdVFlJOAI-gKSlWZrX-P). Further, integration copy number was verified by smFISH (see Methods “smFISH”) to confirm a single integration into one allele of the AAVS1 locus. All primer sequences can be found in Supplementary table 2.

#### ACTB-MS2 and TUBB2B-MS2

WTC AAVS1-MCP-Halo-ACTB-MS2 and WTC AAVS1-MCP-Halo-TUBB2B-MS2 cells, hereafter referred to as “ACTB-MS2” and “TUBB2B-MS2” respectively, were generated by integration of the TV with CRISPR/Cas9 into AAVS1-MCP-Halo with the “SOP: RNP transfection protocol v1.0” from the Allen Institute for Cell Science (https://www.allencell.org/written-protocols.html): sgRNA/Cas9 ribonucleotide protein (RNP) complexes were generated by incubating 1.25 *µ*l (10 *µ*M) of sgRNA (sequence, Synthego) with 2 *µ*g of Cas9 protein for 20 min on ice, then another 25 min at RT while preparing the cells. Cells were nucleofected as described above, the TV was added to the RNP solution just before nucleofection. Cells were plated with 1:1000 CEPT, which was removed the next day, and FACS sorted 8 days after nucleofection to select for cells with stable integration of the construct by constitutive GFP signal. After recovery, cells were treated with different volumes (between 2.5 and 10 *µ*l) of Tat-Cre protein (Merck, scr508) to remove the selection marker. Tat-Cre was washed off the next day. 10 days later, cells were sorted again - this time the GFP negative population was kept. Single cells were directly plated into microgrids with the IsoCell plater (iotaSciences) according to the manufacturer’s protocol (https://iotasciences.com/Technology/), to ensure purely clonal populations. Single clones were grown up and genotyped by PCR with the following primers for the ACTB knock-in: left integration junction: ACTB-KI-LHA_fwd2 and TUBB2B-KI-LHA_rev2; right integration junction: TUBB2B-KI-RHA_fwd2 and ACTB-KI-RHA_rev2; insert/wt band: ACTB-KI-LHA_fwd2 and ACTB-KI-RHA_rev2. Primers used for the TUBB2B knock in were: left integration junction: TUBB2B-KI-LHA_fwd2 and TUBB2B-KI-LHA_rev2; right integration junction: TUBB2B-KI-RHA_fwd2 and TUBB2B-KI-RHA_rev2; insert/wt band: TUBB2B-KI-LHA_fwd2 and TUBB2B-KI-RHA_rev2. PCRs were conducted with the Phusion™ High-Fidelity PCR Master Mix with HF Buffer (NEB, M0531S) according to the manufacturer’s protocol (https://www.neb.com/en/protocols/pcr-protocol-m0530?srsltid=AfmBOorXyKxT6q-Lj-V34jj44p3vIXX1QLATQdVFlJOAI-gKSlWZrX-P). The insert band was cut out of the gel, extracted with the QIAquick Gel Extraction Kit (Qiagen 28704) and subject to Sanger sequencing (primers: ACTB_LHA_FWD and, ACTB_RHA_REV for ACTB-KI, TUBB2B-geno_LHA_fwd and TUBB2B-geno_RHA_rev for TUBB2B-KI) to confirm the seamless and precise integration of the construct. Further, integration copy number was verified by smFISH (see Methods “smFISH”) to confirm a single integration into one allele of the ACTB locus. All primer sequences can be found in Supplementary table 2.

#### ACTB-MS2-NGN2 and WTC-NGN2

To generate the inducible neuron cell lines WTC AAVS1-MCP-Halo-ACTB-MS2-NGN2 (hereafter referred to as “ACTB-MS2-NGN2”) and WTC-NGN2, ACTB-MS2 and WTC wt cells were co-infected with two lentiviruses, containing constructs pLV-EF1a-rtTA-IRES-G418 (Addgene, 84776^64^) and pLV-TetO-hNGN2-hygro (see Methods “Molecular cloning”) as follows: One day after splitting, 1 well of a 6-well plate was incubated with 15 *µ*l of each virus (virus titer was not determined). 2 days later additional 2 ml of media were added without removing the virus. On day 3, confluent wells were split into a 10 cm dish with 1:1000 CEPT and antibiotic selection with G418 (200 *µ*g/ml, InvivoGen ant-gn-1) and hygromycin (1 *µ*g/ml, InvivoGen ant-hg-1) was started on day 4 (with 1:10 000 CEPT). Media was changed every day, on day 9 cells were split again and media was exchanged daily until clear colonies had formed. Single colonies were picked and grown up with antibiotics and without CEPT. To verify successful integration, a subset of the cells of each clone was subjected to 4 *µ*g/ml doxycycline. Clones with close to 100 % neuronal induction efficiency as judged by cell morphology were kept, the others discarded.

### Molecular cloning

#### Targeting vector MCP-Halo

For the AAVS1-MCP-Halo construct, plasmid Puro-Cas9 donor (Addgene, 58409^63^) was used as backbone (BB) and plasmids pCAGGs-MCP-Halo (kindly provided by Jeffrey Chao’s lab, FMI Basel) and pAB1 (see below) were used as templates for PCRs. In total, 4 PCR reactions were set up, 2 to amplify the backbone (in 2 pieces because of the large size), 2 to amplify the inserts. The following primers were used: BB 1: TalenBB_fwd, pIM52_2nd_R; BB 2: TalenBB_rev, pIM52_1st_F; CAGGS: pCAGGs_fwd2, pCAGGs_rev; MCP-Halo: MCP_fwd, MCP_rev2. PCRs were conducted with the NEBNext® High-Fidelity 2X PCR Master Mix (NEB, M0541S) according to the manufacturer’s protocol (https://www.neb.com/en/protocols/pcr-using-nebnext-high-fidelity-2x-pcr-master-mix-m0541srsltid=AfmBOooi3_GOZpIPZKTADD21fxc72sHoKxq4QcX8bp9nUWhnsZTW9hqE). PCR products were purified with the QIAquick PCR purification Kit for PCR Cleanup (Qiagen, 28104). The fragments were assembled with Gibson assembly, using the NEBuilder® HiFi DNA Assembly Master Mix (NEB, E2621S), according to the manufacturer’s protocol (https://www.neb.com/en/protocols/nebuilder-hifi-dna-assembly-reaction-protocol?srsltid=AfmBOorCqQ9sdsct7LxZ1LNAd_Gr9z0BAP1hE8M2XiYKVYJW7gacydqt). As starting material, 50 ng of the largest fragment were used, the amounts of the other fragments were adjusted to result in a 1:1:1:1 ratio. 2 *µ*l of the reaction were transformed into competent DH5α bacteria (Thermo Fisher Scientific, 18263012), colonies were picked the next day and transferred to liquid culture (LB medium with ampicillin). Liquid cultures were minipreped with the QIAprep Spin Miniprep Kit (Qiagen, 27104). Plasmids isolated from single colonies were subjected to restriction digests and the correct sizes of the fragments was confirmed by running them on a 1 % agarose gel (Promega, V3125). Further, the insert region was verified with Sanger sequencing with the following primers: G_stdMCPtm72_R and WPRE_rev. Verified plasmids were expanded in liquid culture and midi-preped with the Zy-moPURE II Plasmid Midiprep Kit (Zymo D4200), including the final optional step to remove endotoxins. All primer sequences can be found in Supplementary table 2.

#### Template MS2 stem loops

To tag endogenous mRNAs with MS2 stem loops, we used a condensed version of the previously published MS2v5 cassette^65^ to facilitate genomic integration. To generate this construct, linker sequences in between individual stem loops of the MS2v5 array were shortened to 3 bp. This yielded a much smaller integration cassette of ∼750 bp (“MS2-3nt”), which is approximately half the size of the previous version, while maintaining identical signalling intensity (data not shown). First, we constructed a vector containing the MS2-3nt stem loops followed by a removable selection marker, which was used as a precursor for the targeting vectors subsequently utilised for endogenous knock-ins via homology-directed repair (pAB1). We generated 3 fragments for Gibson assembly via PCR or restriction enzyme digest: MS2-3nt (template: pIE5 (kindly provided by Jeffrey Chao, FMI Basel); primers: MS2_fwd, MS2_rev), CAGGS-GFP (template: AAV-CAGGs-GFP (Addgene 22212^66^); primers: CAG-GPF_fwd, CAP-GFP_rev) and pBluescript Sk(-) (Addgene 8705^67^; digested with XbaI (NEB, R0145S) and treated with Mung Bean Nuclease (NEB, M0250S) to blunt the ends and prevent re-ligation). PCRs were performed with the NEBNext® High-Fidelity 2X PCR Master Mix (NEB, M0541S), according to the manufacturer’s protocol (https://www.neb.com/en/protocols/pcr-using-nebnext-high-fidelity-2x-pcr-master-mix-m0541srsltid=AfmBOooi3_GOZpIPZKTADD21fxc72sHoKxq4QcX8bp9nUWhnsZTW9hqE). After digestion, pBluescript Sk(-) was run on a 1 % agarose gel and extracted with the QIAquick Gel Extraction Kit (Qiagen 28704). Gibson assembly was done with the NEBuilder® HiFi DNA Assembly Master Mix (NEB, E2621L), according to the manufacturer’s protocol (https://www.neb.com/en/protocols/nebuilder-hifi-dna-assembly-reaction-protocol?srsltid=AfmBOorCqQ9sdsct7LxZ1LNAd_Gr9z0BAP1hE8M2XiYKVYJW7gacydqt). Plasmids were amplified, isolated and verified as described above. All primer sequences can be found in Supplementary table 2.

#### Targeting vector ACTB-MS2 and TUBB2B-MS2

To generate the TV for the ACTB and TUBB2B knock-ins, pAB1 was used as backbone for Golden Gate cloning of the homology arm sequences into the gene of interest. The homology arm sequences (synthesised by GeneWiz, see Supplementary table 2) contain BbsI restriction enzyme cutting sites on either end as well as the sequence of the utilised sgRNA at the 5’ end of the LHA and the 3’ end of the RHA, which results in linearisation of the plasmid inside of the cells. Golden Gate assembly was done by preparing a 15 *µ*l reaction mix containing 10X T4 DNA ligase buffer with 10 mM ATP (1.5 *µ*l, NEB, B0202S), BbsI restriction enzyme (1 *µ*l, ThermoFisher fastDigest, FD1014), T4 DNA ligase (1 *µ*l, NEB, M0202S), vector backbone (1 *µ*l, 100 ng/ul), LHA (1 *µ*l, 50 ng/ul) and RHA (1 *µ*l, 50 ng/ul). The mix was cycled 25 times in a Thermocycler, alternating between 2 min at 37 °C and 5 min at 16 °C, followed by a final digest of unused plasmid for 15 min at 37 °C. The final plasmids were multiplied and verified as described above. Primers for Sanger sequencing were: ACTB: Chim.intr.2_fwd, M13_rev, M13_fwd, CMVenh_rev; TUBB2B: T7, CMVenh_rev, M13_rev. All primer sequences can be found in Supplementary table 2.

#### NGN2 construct for viral infection

pLV-tetO-hNGN2-hygro, hereafter referred to as “NGN2-hygro”, was generated by removing the puromycin resistance cassette from Addgene plasmid 79049^68^ and replacing it with a hygromycin resistance cassette as follows: Addgene plasmid 84776^68^ was used as PCR template for the hygromycin resistance cassette. Primers used to amplify the sequence were HygR2_fwd and HygR_rev and the PCR was performed with the Phusion™ High-Fidelity PCR Master Mix with HF Buffer (NEB, M0531S) according to the manufacturer’s protocol (https://www.neb.com/en/protocols/pcr-protocol-m0530?srsltid=AfmBOorXyKxT6q-Lj-V34jj44p3vIXX1QLATQdVFlJOAI-gKSlWZrX-P). Plasmid 79049 was cut with the restriction enzymes XbaI (NEB, R0145S) and PacI (NEB, R0547S) and the larger band of 9297 bp was extracted from an agarose gel. After digestion of the PCR product with the same enzymes to create compatible overhangs, the backbone and the PCR product (reaction ratio 1:3) were ligated using T4 DNA ligase (NEB, M0202S) and ligase buffer (NEB, B0202S) according to the manufacturer’s protocol (https://www.neb.com/en/protocols/dna-ligation-with-t4-dna-ligase-m0202?srsltid=AfmBOoro7uk9PoawUGR5aySSftQKZq6qomjEVOvVdsPkGx0HoGxj8HyY). The final plasmid was multiplied as described above and verified by full plasmid sequencing. All primer sequences can be found in Supplementary table 2.

All plasmids generated in this study will be made available on Addgene.

### Virus preparation

Lentiviruses (LV) containing either pLVX-EF1a-tetOn-IRES-G418 (Addgene, 8477668) or NGN2-hygro were produced following a four plasmid second generation LV protocol requiring TAT-proteins for LTR dependent integration into the genome. A 1:1:1:1:4 plasmid mix consisting of 1.5 *µ*g of each Gag-pol, Vsvg, Rev and Tat helper plasmids with 6 *µ*g of the respective LV vector was used for transfection into 80-90 % confluent HEK293T cells using TransIT-293 reagent (Mirus Bio, MIR 2705). HEK293T cells were cultured in high glucose DMEM (Gibco, 41965039) supplemented with 10 % (v/v) FBS (Sigma-Aldrich, F2442), 1x Glutamax (Gibco, 35050061) and 0.2 % (v/v) PS (Gibco, 15140122). For LV purification, media was changed to 30 % FBS one day after transfection and collected after 24 hours. A 50x lentiviral suspension was generated in DPBS using Lenti-X™ concentrator (Takara Bio Inc., 631232).

### smFISH

Cells (iPSCs or iNeurons) were seeded onto matrigel-coated coverslips in a 12-well plate at a density of 30 000 cells per well. On the day of harvesting, cells were washed with DPBS (ThermoFisher Scientific 14190144; for neurons specifically DPBS with calcium and magnesium, ThermoFisher Scientific 14040133) and fixed in a 4 % paraformaldehyde (PFA) solution (ThermoFisher Scientific, 043368.9M) for 15 min at RT. The coverslips were washed 3x with nuclease-free PBS and the samples were permeabilised by incubating them in 100 % methanol (Merck, 1060121000) at -20 °C o/n (iPSCs) or in a 0.2 % Triton X-100 (Merck X100-100ML) solution for 10 min at RT (iNeurons). After permeabilisation, cells were washed again 3x with nuclease-free PBS before proceeding with the smFISH protocol. All buffers were prepared on the day of the experiment. Cover slips were incubated in smFISH wash buffer (between 50 % and 65 % formamide (Merck, S4117), depending on the probe set, 2X final concentration of SSC (invitrogen, 15557-036), 200 *µ*M HCl and nuclease-free water) for 5 min (iPSCs) or 10 min (iNeurons). The wash buffer was removed and fresh buffer was added. Incubation was repeated one more time. For hybridisation, coverslips were placed upside down onto 50 ul of hybridisation buffer (50 % dextran sulfate (Merck, S4030), 2X final concentration SSC, formamide at the concentration corresponding to the wash buffer, 1.6 % 1M HCl and nuclease-free water), supplemented with 0.5 *µ*l (12.5 *µ*M) of the corresponding Atto565- or Atto633-labelled probes (all probe sequences can be found in Supplementary table 2). Hybridisation was done o/n at 37 °C in a chamber containing a water reservoir to prevent evaporation of the hybridisation buffer. On the next day, coverslips were washed with wash buffer 3x for 30 min at 37 °C and once with PBS at RT for 5 min before mounting onto microscopy slides by placing them upside down onto a drop of ProLong™ Gold Antifade Mountant with DNA Stain DAPI (invitrogen, P36935). After curing of the Prolong (at least 30 min), coverslips were sealed with transparent nail polish and either directly imaged or stored at +4 °C.

### Immunofluorescence

#### Vascular progenitor cells

Day 5 vascular progenitors were seeded into glass-bottom 4-well dishes (ibidi, 80427). On the day of harvesting, vascular progenitors were fixed o/n at +4°C in a 4 % PFA solution and washed 3x with PBS the next day. Blocking solution consisting of DPBS with 3 % FBS, 1 % BSA (Miltenyi Biotec, 130-091-376), 0.5 % Triton-X (Sigma-Aldrich, 93420), 0.5 % Tween-20 (Sigma-Aldrich, P7949) and 0.01 % sodium deoxycholate solution (Sigma-Aldrich, D6750) was applied for 4 h on an orbital shaker. Primary antibody incubation with mouse-anti-hCD31 (Dako, M0823, 1:100), rabbit-anti-hPDGFR-*β* (Cell Signalling, 3169S, 1:100) and goat-anti-hCollagenIV (Millipore, AB769, 1:100) was done o/n at 4°C on an orbital shaker. The next day, samples were washed 3x with PBS-T (DPBS supplemented with 0.05 % Tween-20) at RT, followed by incubation with secondary antibodies donkey-anti-rabbitAF-568 (ThermoFisher Scientific, A-10042, 1:300), donkey-anti-mouse-AF488 (ThermoFisher Scientific, A-10037, 1:300), donkey-anti-goat-647 (ThermoFisher Scientific, A-21447, 1:300) and DAPI (Sigma-Aldrich, D9542, 1:500) in blocking buffer on an orbital shaker for 5 h at RT. Samples were washed 3x with DPBS and kept in DPBS for imaging.

#### iNeurons

iNeurons were seeded onto matrigel-coated coverslips in a 12-well plate at a density of 30’000 cells per well. On the day of harvesting, they were fixed with 4 % PFA for 15 min and washed 3x with PBS. iNeurons were permeabilised with 0.2 % Triton-X100 in PBS for 10 min and washed 2x with PBS. Samples were blocked with 5 % BSA and 0.2 % TritonX-100 in PBS for 30 min at RT and incubated with primary antibodies chicken-anti-PRPH (ThermoFisher Scientific, PA1-10012, 1:100) and guinea-pig-anti-MAP2 (ThermoFisher Scientific, OSM00149W-100UL, 1:1000) in PBS with 0.1 % Tween and 3 % BSA o/n at +4°C. Samples were washed 2x with 0.1 % Tween and 3 % BSA in PBS for 30 min at RT and secondary antibody incubation with goat-anti-chicken-488 (ThermoFisher Scientific, A-11039, 1:1000) and goat-anti-guinea-pig-AF-647 (ThermoFisher Scientific, A-21450, 1:1000) was done o/n at +4°C. After two washes with PBS the next day, samples were mounted onto imaging slides with ProlongGold + DAPI.

### Sample preparation for live imaging

#### Organoids

Mosaic neural organoids were grown by mixing 5-15 % genetically engineered cells with 85-95 % wt cells. On the day of imaging, neural organoids were incubated with Halo-ligand, coupled to Janelia fluorophore 549 dye (Halo-JF549, final concentration 1 *µ*M), in culture media for 1 hr at 37 °C and washed 3x for 10 min in the incubator to remove excess dye. Organoids were washed in cold HBSS with calcium and magnesium (ThermoFisher Scientific 14025092) by gently pipetting up and down with a cut P1000 tip to remove excess matrigel. Organoids were embedded in 1 % low-melting SeaPlaqueTM Agarose (Lonza, 50101) which was cured for a minimum of 20 min on ice. 200 *µ*m thick live sections were cut with a vibratome (LEICA VT1000 S) at a frequency of 50 Hz and speed of 0.5 mm/s. Slices were laid onto a droplet of Hibernate E media (ThermoFisher Scientific, A1247601), containing ViaFluor® 488 Live Cell Microtubule Stains (1:1000, Biotium 70062) and Vybrant™ DyeCycle™ Violet Stain (1: 1000, ThermoFisher Scientific V35003), in a 35 mm imaging dish (ibidi, 81158). To differentiate between live and dead cells in initial experiments, DRAQ7 (final concentration 3 *µ*M, ThermoFisher Scientific D15105) was added to the mounting media. 4-5 slices were mounted per dish and kept on ice until all samples were processed. To flatten the slices, an 18 mm coverslip (HuberLab, 10.0360.56) was placed on top and samples were allowed to recover at 37 °C for at least 1 hr before imaging on the same day. In case of fixed samples, slices were laid onto a droplet of 4 % PFA and washed 3x with PBS before proceeding with permeabilisation (see Methods “smFISH”).

#### iPSCs, iNeurons and vascular progenitors

Mosaic iNeurons were grown by mixing 30 % genetically engineered cells with 70 % wt cells. iPSCs and vascular progenitors were grown in a non-mosaic fashion. On the day of imaging, cells were incubated with Halo-JF549 and CellMask™ Deep Red Actin Tracking Stain (1:1000, ThermoFisher Scientific A57245) in culture media for 30 min at 37 °C. Dyes were washed out 3x for 5 min in the incubator. For imaging, respective culture medias contained ViaFluor® Live Cell Microtubule Stains (1:1000).

### Image acquisition

Organoid and vascular progenitor imaging was performed on an inverted Nikon Ti2-E Eclipse with a Yokogawa CSU W1 (with Dual T2) spinning disk confocal scanning unit. The microscope was equipped with EMCCD cameras (Andor iXon-Ultra-888) and a SR HP Plan Apo LambdaS 100X silicone oil objective with 0.3 mm working distance (Nikon). For live imaging, temperature and CO2 control were used (37 °C and 5 % CO2). Fields of view (FoVs) were identified in either intact and in-focus tissue regions (neural organoids) or laminin embedding (vascular progenitors) by quickly scanning the sample for labelled cells in the 561 channel at minimal laser power to not bleach the signal before recording. For each FoV, a Z-stack (0.5 *µ*m steps) in all channels with fluorescent signal was acquired, immediately followed by a time series of 100 time points in the Halo dye channel (frame rates: 20 Hz or 2 Hz). iPSCs and iNeurons were imaged in the same setup and with the same acquisition settings, with the exception of using a Plan Apo Lambda 100X immersion oil objective with 0.13 mm working distance (Nikon), instead of the silicone oil objective. All image series of the same sampling frequency were collected from different regions of interest (ROI) across samples. In the majority of the cases, the 2 Hz imaging series was recorded directly after recording the 20 Hz series from the same ROI. For fixed sample imaging, the respective microscope setup was used for the different sample types (silicone oil objective for organoids, immersion oil objective for cells) and Z-stacks (0.2 or 0.5 *µ*m steps) were acquired without temperature and CO_2_ control.

### Small molecule perturbations

#### Translation inhibition with puromycin

Puromycin (Invivogen, ant-pr-1) solution in respective culturing media (2X) was added to the cells directly on the microscope, resulting in a final concentration of 100 *µ*g/ml. Cells were treated for 10 min, then directly imaged.

#### Microtubule depolymerisation with nocodazole

Noco-dazole (Merck, M1404-10MG) solution in respective culturing media (2X) was added to the cells directly on the microscope, resulting in a final concentration of 10 *µ*M. iNeurons were treated for 90 min, vascular progenitors for 45 min, then directly imaged.

#### Actin depolymerisation with cytochalasin D

Cytochalasin D (Selleck Chemicals, S8184) solution in respective culturing media (2X) was added to the cells directly on the microscope, resulting in a final concentration of 10 *µ*M. iNeurons were treated for 15 min, vascular progenitors for 45 min, then directly imaged.

### Neural organoid dissociation

Mosaic week 8 organoids (15 % ACTB-MS2 or TUBB2B-MS2 cells mixed with 85 % WTC wt cells) were dissociated with the Miltenyi Neural Dissociation (Miltenyi, 130-094-802) kit as follows: Two mosaic organoids per cell line were chopped into small pieces with a scalpel and transferred to a 5 ml microcentrifuge tube, where they were washed twice with 1 ml of HBSS without Ca2+ and Mg2+ (Thermo Scientific, 14175095). After removing the supernatant, 1 ml of enzyme mix 1 (Enzyme P/Papain + Buffer X from Miltenyi Neural dissociation kit) was added to each sample and incubated at 37°C in the water bath or inside the incubator for 15 min. Following the first incubation step, 15 *µ*l of enzyme mix 2 (Enzyme A/DNase + Buffer Y from Miltenyi Neural Dissociation kit) were added and cells were triturated 5–10 x with a p1000 pipette and triturated again 5–10 x with a p200 before another 15 min of incubation at 37°C. Samples were then further triturated with a p200 pipette and incubated for another 10 min at 37°C. If cells were still clumping together, samples were triturated once more with a p200 pipette. Quality of the dissociation was monitored by observing 1 *µ*l of cell suspension under the microscope at a magnification of 10 X. If clumps persisted, further 5 min incubation and trituration steps were performed. When dissociation was complete, cells were filtered and centrifuged for 5 min at 300 g using a swing centrifuge. The supernatant was removed and cells were frozen in 50 % CryoStor (Merck, C2874) until further processing.

### Single-cell transcriptome library generation and sequencing

Single-cell transcriptome cDNA libraries from thawed cell suspensions were generated as described previously.^69^ Briefly, the Chromium Next GEM Single Cell 3^*′*^ v3.1 kit (10x Genomics) was used and library preparation was done according to the 10x protocol (CG000204 Rev D for v3.1). 10X libraries were sequenced in an S1 flow cell (Illumina NovaSeq technology) by the Genomics Facility Basel, using a paired-end 28/10/10/90 configuration (Read1/IDX i7/IDX i5/Read2). Cell Ranger (v3.1.0, 10X Genomics) with reference genome GRCh38-3.0.0 was used to derive gene-cell count matrices from the sequencing read (fastq) files of the gene expression library.

### Image processing and feature extraction

#### Cell segmentation

##### Neural organoids

To segment single MCP-Halo-labelled cells in organoids, binary cell masks (whole cells and nuclei) were manually curated using the “Freehand selection” tool in Fiji.^70^ To create cytoplasm masks, nuclei masks were subtracted from whole cell masks.

##### iNeurons

To segment single MCP-Halo-labelled cells in iNeuron culture, cytoplasm and nucleus masks from two experimental batches were hand-corrected and combined by expanding and assigning each nucleus to its overlapping cell label, followed by hole-filling and removal of small segments. A Cellpose model (v3.1.1.1)^71^ was fine-tuned from the pre-trained cyto3 base on single-channel z-projection images using the RAdam optimiser (learning rate = 0.005) for 2,000 epochs. For inference, the fine-tuned model was used with a flow threshold of 0.4, and a cell probability threshold of 0.0. Nuclei masks were created with Ilastik^72^ and subtracted from the whole cell masks. As a final step, all cytoplasm masks were manually corrected in Napari.^73^

##### Vascular progenitors and iPSCs

Cell segmentation was performed using intensity-based thresholding in the MCP-Halo channel. For nuclear and whole cell segmentation, a global Otsu^74^ threshold was calculated respectively from maximum intensity projections and scaled empirically. Binary cell masks were generated by thresholding, followed by hole filling, small object removal, and Gaussian smoothing to regularise boundaries. Individual cells were separated using marker-controlled watershed segmentation. Labeled nuclei were used as markers, and the watershed was applied to the negative distance transform within the whole-cell mask to resolve overlapping cells. To create cytoplasm masks, nuclei masks were subtracted from whole cell masks. All vascular progenitor cytoplasm masks were manually corrected in Napari.^73^

#### Denoising, spot detection and tracking

Raw acquired time series images (TYX) were denoised with Noise2Void2 (N2V2),^75,76^ a self-supervised content-aware image restoration technique. Two-channel time series images were used for spot detection, where spot identification was performed on the denoised channel and spot intensities were extracted from the raw channel at the determined spot coordinates. Masked images were created by applying the corresponding label images to all T slices of each two-channel series. For spot detection, masked images were subjected to a combination of the Laplacian of Gaussian (LoG) filter and h-maxima^77^ thresholding. The LoG sigma is derived from the theoretical Abbe resolution limit based on the emission wavelength maximum (570 nm for dye JF549) and numerical aperture of the objective (NA=1.35 for organoid images; NA=1.41 for cell images). The h-maxima threshold is chosen to be the standard deviation of the mean intensity of the regions of interest of each image multiplied by an empirically selected factor, depending on the tissue type and age of the sample. This results in a unique per-image threshold that compensates for differences in illumination at varying z-positions and enables the application of a single multiplication factor for all images of a given batch of experiments. To identify a suitable threshold for each sample kind and age, different factors between 3 and 6 were tested. The results were visualised by laying the detected spots over a representative sample of images and the factor which captured most spots without detecting stochastic increases in background intensities was chosen for the respective sample kind. The detected spots were refined to sub-pixel coordinates by fitting a 2D Gaussian. When available, intensities of the live actin and tubulin stains were extracted for every detected spot coordinate pair and included in the output table. Tracking was done with the basic Trackpy linking function^78^, with the following parameters: linking distance of 5 or 7 pixels (20Hz or 2Hz regime, respectively), 2 allowed gaps and a minimum track length of 5.

### Mobility features

### Sliding diffusion coefficient

To identify sub-track behaviours, we estimated the diffusion coefficient, derived from mean squared displacement calculations, using a sliding window approach as follows: Each track was divided into sub-tracks with a sliding window size of 5 and a step size of 1. Within each window, time-averaged mean squared displacements (TAMSD) were computed for each time lag. The diffusion coefficient per window (sliding diffusion coefficient) was derived by fitting a linear model to the mean squared displacements of the first three time lags to ensure each point used for fitting was derived from an average of at least three displacement measurements within the five-frame window. The extracted slope values were converted to diffusion coefficients using the relation for 2D diffusion:

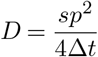

where *s* is the slope, *p* is the pixel size (0.134 *µ*m) and Δ*t* is the time interval between frames (0.05 s (20 Hz) or 0.5 s (2 Hz)).

#### Diffusion coefficient and anomalous exponent $*α*

To quantify the diffusion coefficients and anomalous diffusion behavior of each identified sub-track and whole track, we computed the TAMSD along the entire (sub-)trajectory. The diffusion coefficient and anomalous exponent *α* were estimated by fitting linear models to the mean squared displacement as a function of time lag in linear scale and log–log scale, respectively. Only the first 20 % of available lag points (with a minimum of five) were used for fitting, ensuring slope estimation was done in the linear range of the curve and while maintaining statistical robustness. The extracted slope values were converted to diffusion coefficients as described above.

#### Directionality

Directionality of each identified sub-track was quantified as the ratio of the net displacement to the total path length. For each track, the Euclidean distance between the start and end points was divided by the sum of distances between consecutive positions along the trajectory. This ratio ranges from 0 to 1, where 1 indicates perfectly straight motion and values closer to 0 indicate more convoluted or random movement.

### Intensity features

#### Cytoskeleton intensity

To obtain actin and tubulin fluorescence intensities across trajectories as a proxy for co-localisation of single particles with the cytoskeleton, per-spot cytoskeleton dye intensities were normalised using z-score normalisation within each image. Normalised intensities were stored as trajectory-averaged values.

### Distance features

#### Distance to nucleus

To quantify the distance of single particles to the nucleus, two metrics were computed for each trajectory. First, the mean distance to the nucleus was calculated by averaging the per-frame distance-to-nucleus values across each individual track. Second, the net change in nuclear distance was obtained by subtracting the distance at the first time point from that at the final time point within each trajectory.

#### Distance to edge

To quantify the distance of single particles to the cell boundary, two metrics were computed per trajectory. First, the mean distance to the cell edge was determined by averaging the per-frame distance-to-edge values across all time points within each trajectory. Second, the net change in distance to the edge was calculated by subtracting the distance at the initial time point from that at the final time point within each trajectory.

### Cell compartment masks

To generate whole-cell masks, nucleus and cytoplasm segmentations were combined as follows: For each nucleus, the nuclear mask was iteratively expanded by binary dilation (1 pixel per iteration, up to a maximum of 20 pixels) until the expanded region fully overlapped with the corresponding cytoplasmic mask. Expanded nuclei were then merged with cytoplasmic masks to obtain complete single-cell masks. To classify subcellular compartments, euclidean distance transforms were computed for each cell and spatially smoothed using a uniform filter to obtain a continuous measure of distance from the cell boundary. Regions with large smoothed distances were classified as soma-outgrowth. Intermediate-distance regions were classified as neurites. Small disconnected objects were removed to exclude segmentation noise. Single-molecule tracks were assigned to cellular compartments based on their spatial position within the compartment masks. For each localisation, the corresponding compartment label was extracted from the mask at the spot’s (x,y) coordinates. To distinguish soma from outgrowth (the latter including all non-elongated structures distant from the nucleus, such as branch points and growth cones), localisations within the soma–outgrowth mask were further classified using the distance to the nucleus: positions closer than 5 *µ*m to the nucleus were assigned to the soma, while more distal positions were assigned to outgrowths, finally splitting each cell into three subcellular compartments: “soma”, “outgrowth” and “neurite”.

### Hidden Markov Modelling

To identify hidden states underlying observed particle trajectories, Hidden Markov modelling was performed on per-track feature vectors containing the sliding diffusion coefficient values of each track, ordered by sliding window. The model was trained on a 1 % random sub-sample across all biological model systems and perturbations to obtain a standardised framework as follows: A two-step refinement of Gaussian Hidden Markov Models (Gaussian HMMs), implemented from the hmmlearn library (https://hmmlearn.readthedocs.io/en/latest/index.html), was performed. First, a 3-state Gaussian HMM with full co-variance matrices was fitted to the input features using 100 random initialisations. The best-performing model, selected based on the log-likelihood score, was used to assign initial state labels. Subsequently, the state associated with the lowest mean sliding diffusion coefficient, i.e. the least mobile state, was identified and isolated for further refinement. A second 2-state Gaussian HMM was then fitted to the subset of data corresponding to this least mobile state. The resulting substates were relabeled such that the majority substate retained the original label, while the minority was assigned a new state identifier, resulting in a total of four states (states 0, 1, 2, 3). Using the trained model, hidden states were predicted for all sliding windows of all trajectories across the dataset. Each x-y coordinate pair was assigned the hidden state corresponding to the sliding window centered on that point. Consecutive sequences of the same state within a trajectory were grouped into unique sub-tracks, with a minimum required length of three frames. Shorter segments (i.e., fewer than three consecutive points) were reassigned to adjacent tracklets. Mobility features diffusion coefficient, *α*, instantaneous diffusion coefficient and directionality were calculated for all unique sub-tracks as described above. To capture episodes of directed movement, a fifth state was introduced: Any sequence within a track with an *α* value greater than 1.7 and a directionality value greater than 0.85 (thresholds were determined empirically) was assigned a new hidden state (state 4). To avoid inflating the fraction of directed particles with short, transient high-*α*, high-directionality segments, only sequences that spanned at least eight frames were considered.

### Track clustering

#### Track encoding

For each track, di-grams representing transitions between hidden states were generated. Tracks consisting of a single state were assigned a self-transition di-gram (e.g., 0→0, 1→1, 2→2, 3→3, or 4→4). Each unique digram was weighted by its frequency within the track, and the weights were normalised to sum to 1 to account for differences in track length. This procedure yielded a unique feature vector per track, encoding the relative frequency of all possible state transitions.

#### Generation of pseudo-tracks

A k-nearest-neighbor (kNN) graph was constructed from the above mentioned feature matrix using the Jaccard distance metric with k+1 neighbors, after which self-neighbors were removed. Distances and neighbor indices were obtained with the NearestNeighbors implementation from scikit-learn^79^, and edge weights were defined as exp(d), where d represents the Jaccard distance. The weighted graph was represented in igraph^80^, and Leiden clustering was performed using the leidenalg package (https://pypi.org/project/leidenalg/) with the RBConfigurationVertexPartition, using 15 nearest neighbours and a specified resolution parameter (5 for 20 Hz regime, 3 for 2 Hz regime). Clustering resulted in 175 unique clusters (20 Hz regime) and 98 unique clusters (2 Hz regime) that were termed pseudo-tracks, as they each summarised one particular type of transcript motion.

#### Hierarchical clustering

For each pseudo-track, a feature matrix based on weighted di-gram occurrences was constructed in an equivalent way to the feature matrix created for single tracks. Hierarchical clustering was performed on the obtained feature matrix after standardisation to zero mean and unit variance using the scipy.cluster.hierarchy module from the scipy library^81^. A linkage matrix was computed using the Ward method with Euclidean distance, and cluster labels were obtained by cutting the dendrogram to yield a predetermined, empirically tested number of 15 clusters (20 Hz regime) and 12 clusters (2Hz regime).

### Downstream calculations

#### Subsetting

For each specific analysis, pseudo-tracks and clusters were subset to include only tracks corresponding to the relevant conditions. Pseudo-tracks and clusters containing fewer than 10 entries after subsetting were excluded, as such small groups were considered insufficient for reliable conclusions. Node sizes in the circular lineage-like plots were scaled according to the number of particles within each subset cluster.

#### Fraction creation

To correct for differences in dataset size across conditions, the track count for each condition–cluster pair was normalised by the total number of particles belonging to that condition across all clusters as follows:

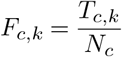

*T*_*c,k*_ = no. of tracks for condition *c* in cluster *k*

*N*_*c*_ = total no. of particles for condition *c* across all clusters

This yielded a balanced fraction of each condition within every cluster, which was then used for downstream analyses.

#### Enrichment score

For each pseudo-track and each cluster, the relative enrichment of each condition was computed by comparing the within-pseudo-track or within-cluster fraction of each condition to the average fraction of that respective condition across all other groups or clusters. Specifically, for each group or cluster, the fraction of observations assigned to each condition was divided by the mean fraction of that condition in the remaining groups or clusters, respectively. Extreme values were clipped using the 5^th^ and 95^th^ quantiles to limit the influence of outliers.

#### Log_2_ fold-change

For each pseudo-group and Leiden cluster, the log_2_ fold-change of the perturbation relative to the unperturbed condition was computed. Observations were aggregated per group or cluster to form contingency tables of perturbation counts. The log_2_ fold-change was calculated as the ratio of the perturbation frequency to the frequency in the unperturbed condition, with a small pseudocount added to avoid division by zero.

### Single-cell RNA sequencing analysis

Single-cell RNA-sequencing data were processed using Scanpy.^82^ Count matrices from Cell Ranger were imported from filtered feature-barcode matrices, and gene and cell identifiers were made unique. Quality control metrics were computed per cell, including total counts, number of detected genes, and the fraction of mitochondrial, ribosomal, and hemoglobin transcripts. Cells with fewer than 100 detected genes and genes expressed in fewer than three cells were removed. Doublets were identified using Scrublet.^83^ Raw counts were normalised to total counts per cell and log-transformed. Highly variable genes were identified (top 2,000), and principal component analysis was performed on this feature set. A k-nearest-neighbor graph was constructed in PCA space and used to generate a UMAP embedding for downstream visualisation.

## Code availability

All image processing and feature extraction code is available on our public GitHub repository: https://github.com/fmi-basel/singlemolecule-tracking All data processing and analysis code is available on our public GitHub repository: https://github.com/quadbio/trackline

## ACKNOWLEDGEMENTS

We would like to thank Jeffrey Chao (FMI Basel) for productive discussions and generous sharing of resources. We would like to thank all members of the Treutlein, Voigt and Chao labs for discussions and feedback. We would like to thank the imaging facilities of the FMI (Facility of Advanced Imaging and Microscopy) and the UZH (Center for Microscopy and Image Analysis) for their support, as well as Claudia Brunner-Barrios for help with experiments and George Hausmann and Leonardo Maffioli for proof-reading the manuscript.

## AUTHOR CONTRIBUTIONS

A-D.B. performed all experiments, with support from R.O., M. Seimiya, M. Santel. and F.V.; A-D.B. performed all image processing with help from T-O.B.; A-D.B. performed all image feature extraction and downstream analyses with help from G.G.; A-D.B. analysed single-cell transcriptome data; T-O.B. wrote the spot detection algorithm and helped with developing the custom single-particle tracking pipeline; G.G. helped with image feature extraction and Hidden Markov modelling; R.O. helped with cell and organoid culture; M. Seimiya cultured and performed immunofluorescence experiments in blood vessel organoids; M. Santel helped with generating single-cell transcriptome data; F.V. performed Northern Blots and immunofluorescence experiments in iNeurons; A-D.B., B.T. and F.V. designed the study and wrote the manuscript.

## Supplementary figures

**Figure S1.**
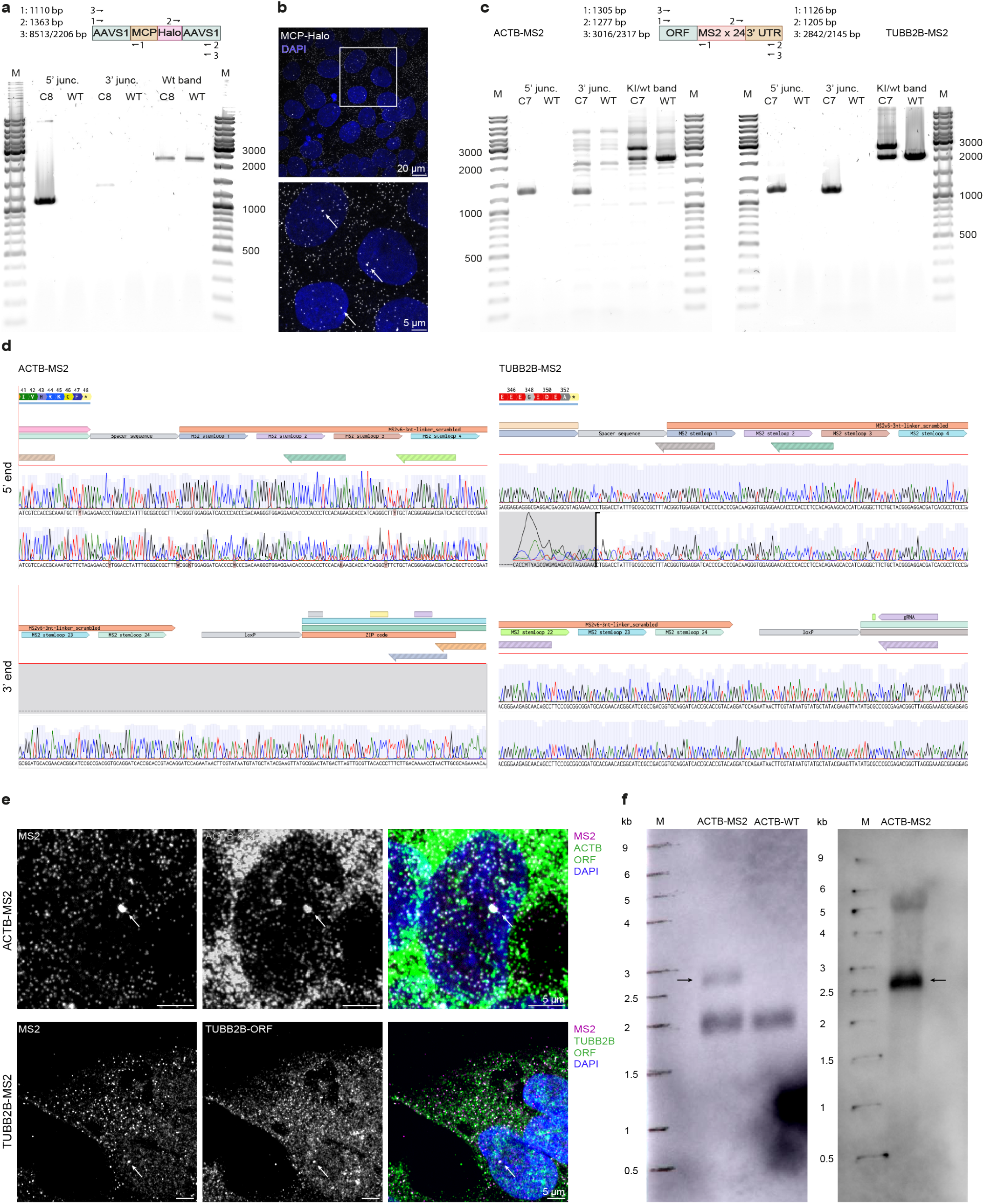
Integration of the MS2/MCP system in human iPSCs. **a**. PCR analysis of the 5^*′*^ junction, 3^*′*^ junction, and full MCP-Halo insert at the AAVS1 locus in a selected clone. Expected band sizes: 5’ junc., 1110 bp; 3’ junc., 1363 bp; insert, 8513 bp; wild type, 2206 bp. Junction PCRs demonstrate the successful integration of the MCP-Halo construct. The full insert could likely not be amplified due to its large size and consequent preferential amplification of the shorter PCR product that spans the unedited AAVS1 locus. The presence of the latter confirms heterozygous insertion. **b**. Single-molecule FISH against MCP-Halo. One transcription site per cell (white arrows) is observed, confirming a single heterozygous insertion of the construct. **c**. PCR analysis of the 5^*′*^ integration junction, 3^*′*^ integration junction, and full MS2 cassette confirms successful integration of the MS2 stem loop array into the desired endogenous loci of either ACTB (left) or TUBB2B (right). Selected single clones are shown for both cell lines. Expected band sizes for ACTB: 5’ junc., 1305 bp; 3’ junc., 1277 bp; insert, 3016 bp; wild type, 2317 bp. Expected band sizes for TUBB2B: 5’ junc., 1126 bp; 3’ junc., 1205 bp; insert, 2842 bp; wild type, 2145 bp. The presence of a wild type band in the edited cells demonstrates heterozygosity of the MS2 insertion for both cell lines. **d**. Sanger sequencing of isolated full length PCR product confirms seamless integration of the MS2 cassette into the endogenous ACTB (left) and TUBB2B (right) loci. **e**. Single-molecule FISH against MS2 stem loops and either the ACTB (top) or TUBB2B (bottom) open reading frame (ORF). A single double-labelled transcription site (white in overlay, indicated by arrows) is detected in the nucleus, confirming heterozygous insertion of the construct, while excluding additional random integrations into the genome. **f**. Left: Northern blot analysis of ACTB-MS2 cells using a probe targeting the ACTB ORF. Expected band sizes: wt transcript: 1800 nt; MS2-tagged transcript: 2500 nt. The black arrow indicates the MS2-tagged transcript. Right: Northern blot analysis of ACTB-MS2 cells using a probe targeting the MS2 stem loops. Expected band size: MS2-tagged transcript: 2500 nt. The black arrow indicates the MS2-tagged transcript. The smeared-out band roughly double in size likely corresponds to an oligomerised running artefact. No additional bands were detected, confirming the integrity of the tagged transcript.

**Figure S2.**
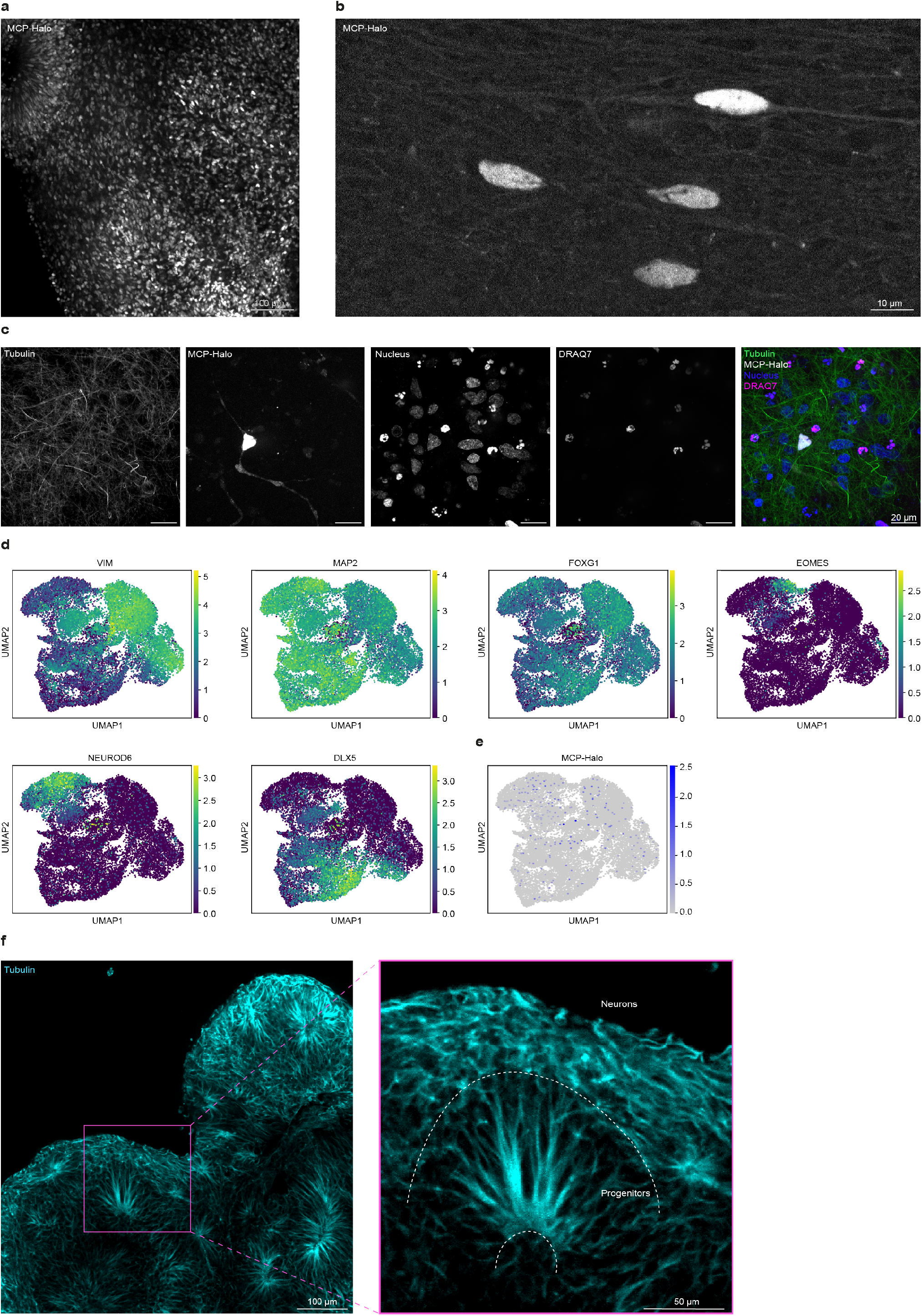
Organoid generation for live single-molecule imaging. **a**. Maximum-intensity projection of a week 12 neural organoid generated from MCP-Halo–expressing cells. **b**. High magnification image of a week 12 MCP-Halo organoid showing the absence of detectable single-molecule spots. **c**. Tissue section of a week 8 neural organoid stained with live tubulin and a nuclear dye, together with a live/dead stain (DRAQ7) to assess tissue integrity following slicing. An expected number of dead cells is observed with nuclear morphology clearly distinct from healthy cells. **d**. Gene expression feature plots from combined single-cell RNA-sequencing data of week 8 organoids generated from mosaic cultures containing either 15 % ACTB-MS2 or TUBB2B-MS2 cells and 85 % wt cells. Colors indicate log1p-transformed, library size–normalised expression (counts per 10’000). As expected, most cells were positive for MAP2, confirming their neuronal identity, with a sub-population of cells strongly expressing the neuronal progenitor marker VIM. Most cells were further FOXG1 positive, identifying them as belonging to the developing cerebral cortex. Of the MAP2 and FOXG1 positive cells, a sub-population expressed EOMES, demonstrating the presence of intermediate progenitor cells. Two other sub-populations expressed NEUROD6 and DLX5 in a mutually exclusive manner, confirming that cells of both the dorsal and ventral telencephalon lineage were present. **e**. Cells expressing the MCP-Halo construct (and by extension MS2-tagged transcripts) are highlighted in blue. Notably, the number of cells with detectable MCP-Halo levels was relatively low (1 %) compared to their input fraction (15 %), suggesting that genetically-engineered cells were largely outcompeted by their wt neighbours. **f**. Live tubulin staining outlining the structure of a week 5 neural organoid. The neuronal layer gets established perpendicular to the progenitor zone. Distinct regions in the organoid are indicated with dashed lines. For all time points, cells embedded in radially aligned tissue were termed “progenitors”. In week 5 organoids, cells in regions outside of progenitor zones were termed “W5 neurons” while in week 8 organoids, these cells were termed “W8 neurons”.

**Figure S3.**
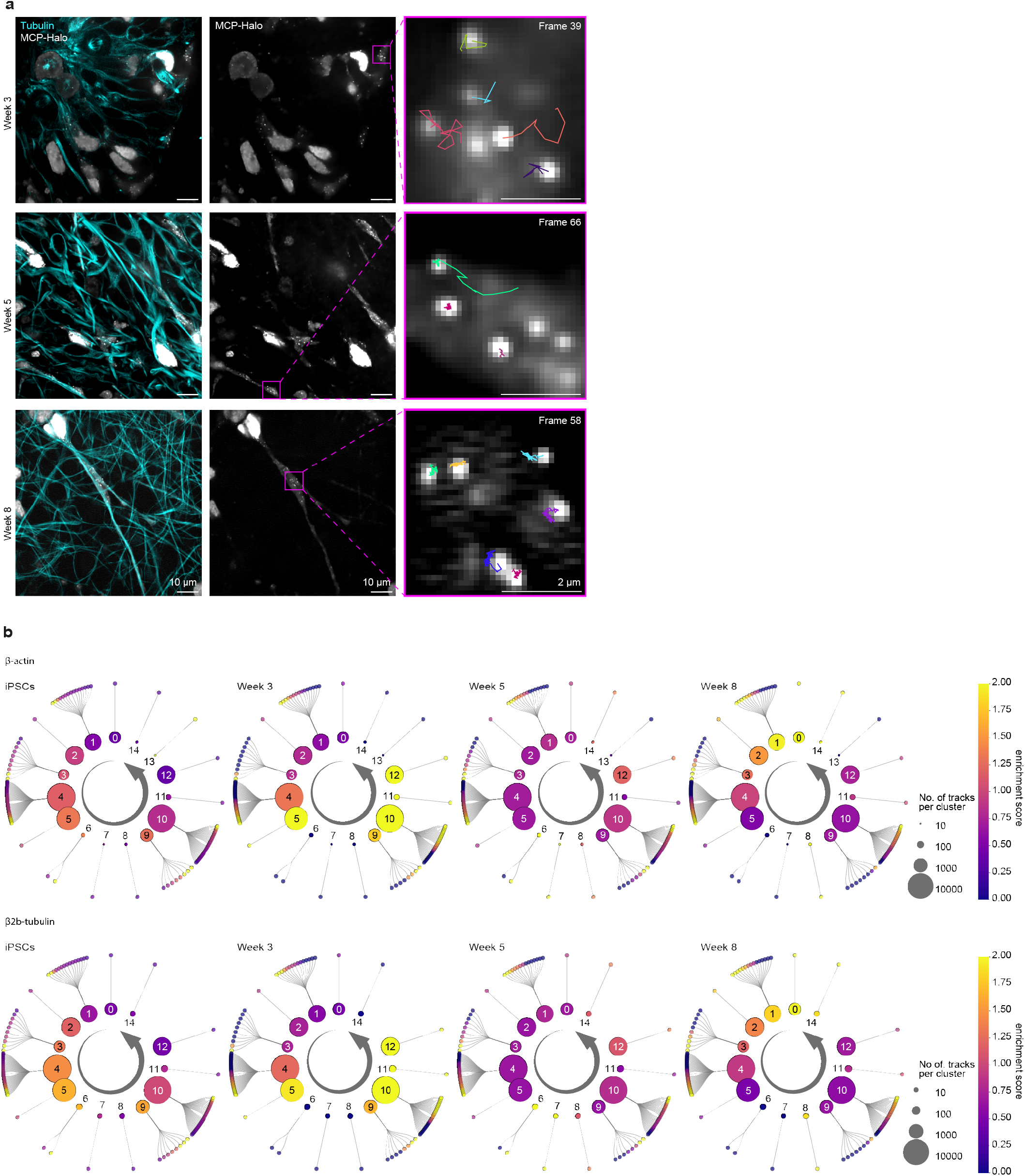
Live imaging of *β*2b-tubulin mRNA in neural organoid slices shows similar shifts in mobility cluster composition as for *β*-actin transcripts. **a**. Live imaging of tissue sections at three developmental time points. TUBB2B-MS2–labelled cells interspersed with wt cells are outlined in white. Insets show individual molecules and their corresponding tracks. **b**. Mobility cluster and pseudo-track enrichment for tagged *β*-actin (top) and *β*2b-tubulin (bottom) transcripts across developmental time points. Enrichment was calculated as the fraction of particles in a given cluster or pseudo-track at one time point divided by the average fraction across all time points. While no clusters appeared enriched in iPSCs, most sub-diffusive and diffusive clusters (5 and 9-12, respectively) were highly enriched by week 3 in both *β*-actin- and *β*2b-tubulin-MS2 organoids. This effect was ameliorated by week 5 when only clusters 6 and 7 were enriched, both of which contain a stalled, sub-diffusive and diffusive component, suggesting increased switching between mobile and tethered phases. By week 8, the effect was completely inverted, with clusters consisting of exclusively stalled and constrained particles (clusters 0-2) as well as cluster 8 (equally consisting of stalled and diffusive states) being strongly enriched. Interestingly, clusters 13 and 14 are specifically enriched at distinct time points, even though they are both dominated by directed behavior: While cluster 13 containing both a directed and a sub-diffusive component (only present for *β*-actin transcripts) is strongly enriched in iPSCs, cluster 14 consisting purely of directed particles is most strongly enriched in week 8 organoids.

**Figure S4.**
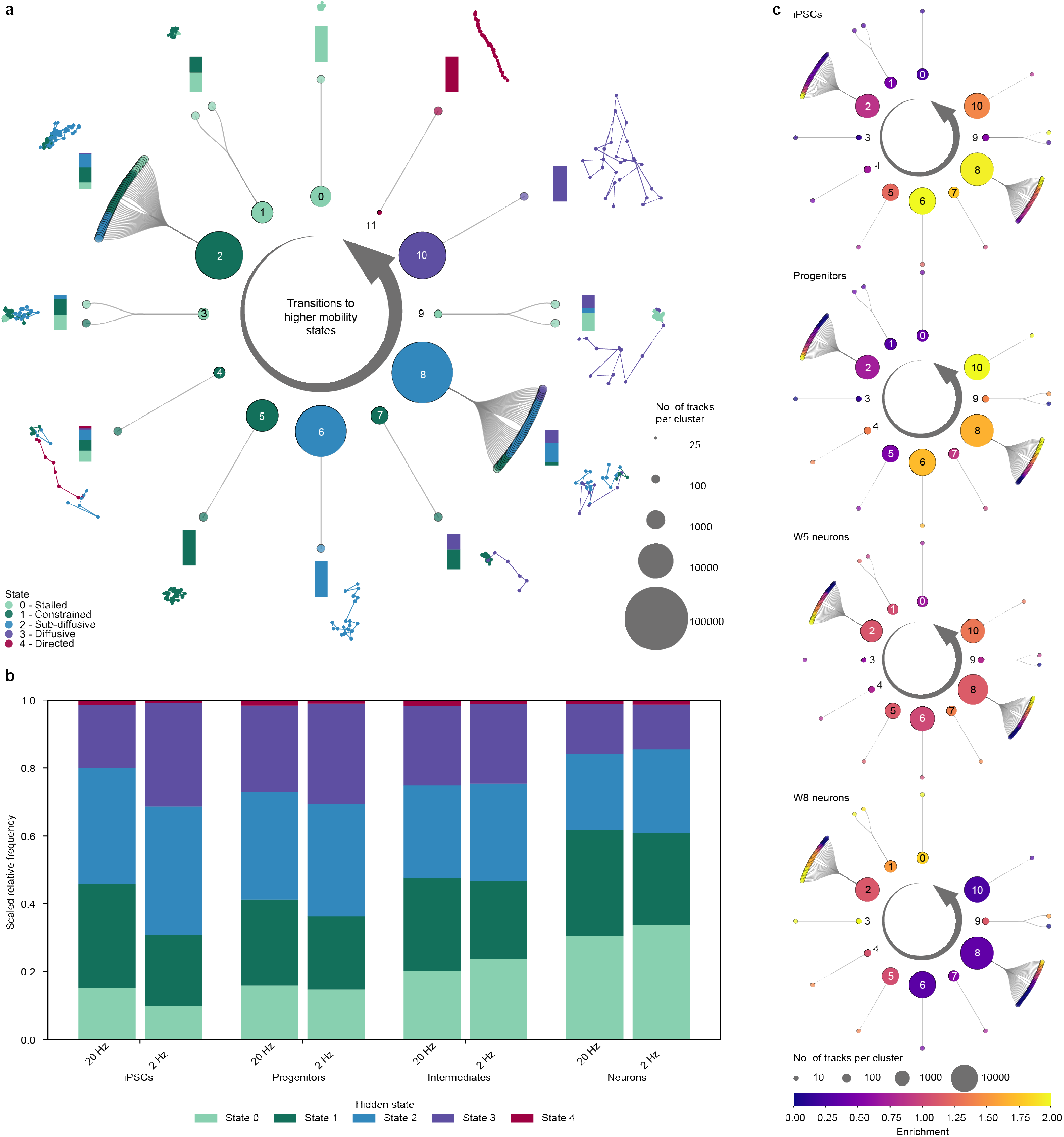
2 Hz imaging regime of *β*-actin transcripts in neural organoids confirms stable fraction of stalled and constrained particles in neurons. **a**. Lineage-tree-like representation of mobility classes, data acquired at 2 Hz. Inner ring: Top-level clusters ordered by increasing di-grams (transitions to higher-mobility states) and coloured by the most frequent hidden state; node size indicates the number of tracks per cluster. Outer ring: Pseudo-tracks assigned to each mobility cluster, coloured by their most frequent hidden state. Stacked bars denote the fraction of hidden states per cluster. Representative tracks are shown. **b**. Hidden-state distributions of tagged *β*-actin transcripts imaged at 20 Hz vs. 2 Hz in iPSCs and across cell types in neural organoids. Min–max-normalised square-root–transformed fractions are shown. **c**. Mobility cluster and pseudo-track enrichment for tagged *β*-actin transcripts across cell types. Enrichment was calculated as the fraction of particles in a given cluster or pseudo-track within one cell type divided by the average fraction across all cell types. In accordance with the 20 Hz regime, the most diffusive cluster (10) is strongly enriched in progenitors, no particular enrichment is observed in W5 neurons, and moderate to strong enrichment of clusters containing the stalled state (cluster 9 and clusters 0, 1, 3, respectively) is present in W8 neurons. This confirms that the observed cell-type-dependent particle mobility phenotype is robust across the two different time scales.

**Figure S5.**
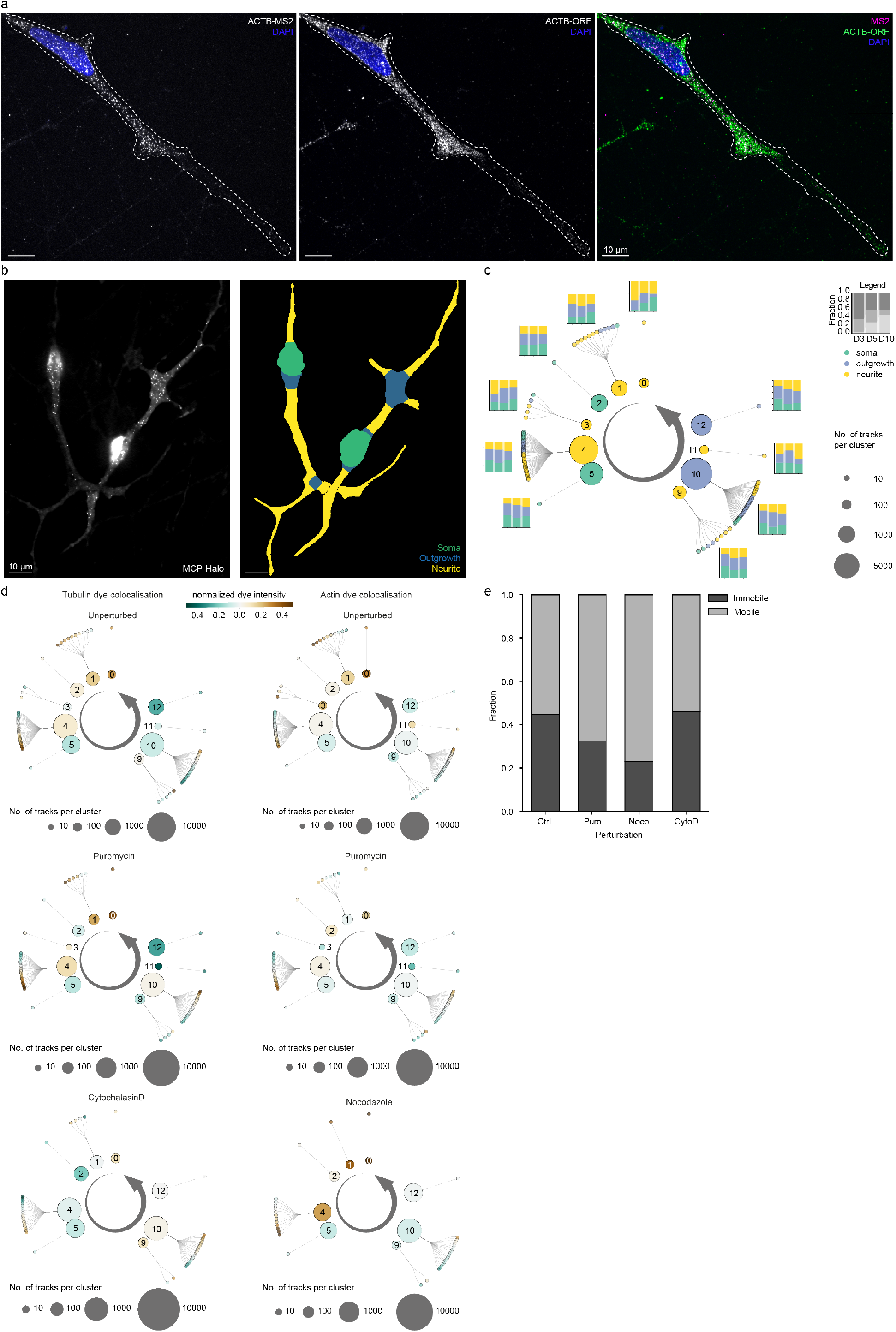
Subcellular compartment- and cytoskeleton-specific targeting of *β*-actin mRNA in iNeurons. **a**. smFISH in day 10 iNeurons showing co-localisation of ACTB-MS2 and ACTB-ORF signals (white in overlay). **b**. Left: Single-frame image of day 5 iNeurons. Right: Cell-compartment mask generated from combined cell and nuclear masks (see Methods). **c**. Lineage-tree-like representation of the dominant cell compartment per cluster (inner ring) or pseudo-track (outer ring) across cumulative iNeuron data. Bar plots surrounding the circle show compartment fractions per cluster resolved by developmental time point. **d**. Z-score–normalised cytoskeletal intensities per track, averaged by cluster (inner ring) and pseudo-track (outer ring) and resolved by perturbation in day 5 iNeurons. For each detected spot ((x,y) coordinate pair), intensities from live-dye channels were extracted, normalised, and averaged per track, providing a proxy for mRNA co-localisation with the actin or tubulin cytoskeleton (see Methods). **e**. Fraction of immobile vs. mobile particles (aggregation of states 0 and 1, and states 2 and 3, respectively) in unperturbed and perturbed conditions in day 5 iNeurons.

**Figure S6.**
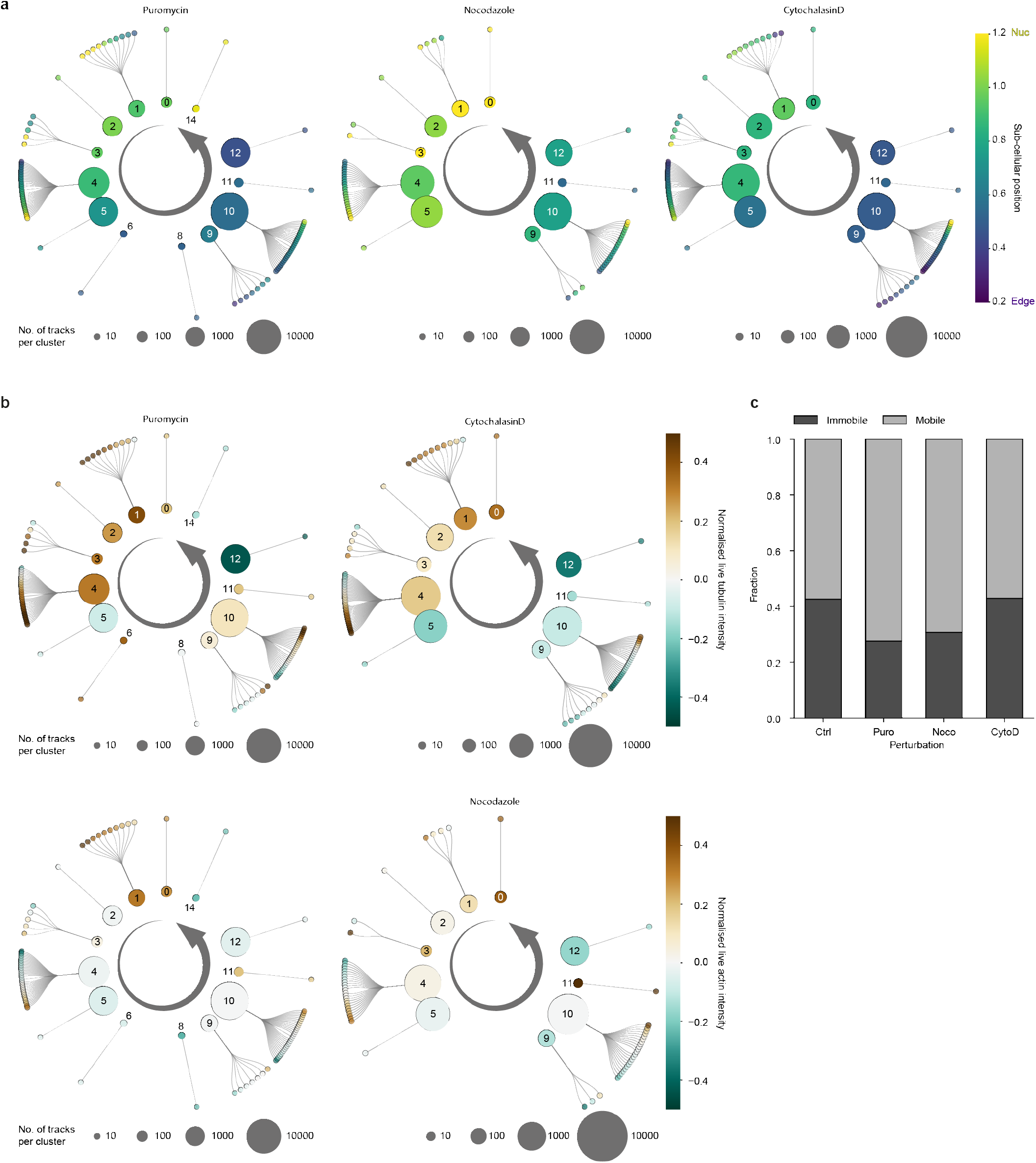
Subcellular position and association to cytoskeletal components of *β*-actin transcripts in migrating vascular progenitor cells. **a**. Distance to edge/nucleus ratio averaged by cluster (inner ring) and pseudo-track (outer ring) in day 10 sprouting cells and resolved by perturbation. Higher values indicate proximity to the nucleus, whereas lower values indicate proximity to the cell edge. **b**. Z-score–normalised cytoskeletal intensities per track, averaged by cluster (inner ring) and pseudo-track (outer ring) in day 10 sprouting cells and resolved by perturbation. For each detected spot ((x,y) coordinate pair), intensities from live-dye channels were extracted, normalised, and averaged per track, providing a proxy for mRNA co-localisation with the actin or tubulin cytoskeleton (see Methods). **c**.Fraction of immobile vs. mobile particles (aggregation of states 0 and 1, and states 2 and 3, respectively) in unperturbed and perturbed conditions.

## Supplementary tables

***Supplementary table 1***. Summary of number of particles tracked across cell lines, model systems and ROIs.

***Supplementary table 2***. Summary of primers, smFISH probes and gene fragments used for this study.

